# Human-caused wolf mortality persists for years after discontinuation of hunting

**DOI:** 10.1101/2022.12.20.521126

**Authors:** Roman Teo Oliynyk

## Abstract

By the mid-20th century, wolves were nearly extinct in the lower 48 states, with a small number surviving in northern Minnesota. After wolves were placed on the endangered species list in 1973, the northern Minnesota wolf population increased and stabilized by the early 2000s. A wolf trophy hunt was introduced in 2012–2014 and then halted by a court order in December 2014. The Minnesota Department of Natural Resources collected wolf radiotelemetry data for the years 2004–2019. Statistical analysis showed that wolf mortality remained close to constant from 2004 until the initiation of the hunt, and that mortality doubled with the initiation of the first hunting and trapping season in 2012, remaining at a nearly constant elevated level through 2019. Notably, average annual wolf mortality increased from 21.7% before wolf hunting seasons (10.0% by human causes and 11.7% natural causes) to 43.4% (35.8% by human causes and 7.6% natural causes). The fine-grained statistical trend implies that human-caused mortality increased sharply during the hunting seasons, while natural mortality initially dropped. While gradually diminishing after the discontinuation of the hunt, human-caused mortality remained high, which could indicate a lasting shift in human attitudes and post-hunt changes in wolf behavior and wolf pack composition.

## Introduction

By the mid-20th century, wolves were nearly extinct in the lower 48 states of the United States, with only 450–700 wolves remaining in northern Minnesota in the 1950s [1, 2]. After wolves were protected by the federal Endangered Species Act of 1973, the northern Minnesota wolf population gradually increased and stabilized in the early 2000s [3]. In late December 2011, the US Fish and Wildlife Service removed wolves in the western Great Lakes region from the federal endangered species list [4]. The following year, Minnesota and other states established a wolf trophy hunting and trapping season, which was extended in 2013 and 2014 [5–8]. In December 2014, a federal judge placed wolves back on the endangered species list, thereby halting further hunting and trapping in Minnesota [9].

According to wolf population surveys conducted by the Minnesota Department of Natural Resources (MN DNR) [3, 10, 11], wolf population estimates for the years 2012–2014 varied from approximately 2,200 to 2,400. MN DNR *Minnesota Wolf Season Reports* [6–8] accounted for 413 wolves killed in 2012, 238 in 2013, and 264 in 2014. Furthermore, the MN DNR collected radiotelemetry data on wolf movements and mortality between 2004 and 2019; see Chakrabarti et al. [12] for a description of the telemetry data collection methods. In this paper, “hunting and trapping season” will often be abbreviated to “hunting season”—inclusive of trapping—for conciseness.

This study aimed to compare wolf survival and mortality for two specific time ranges: before and after the initiation of the first wolf hunting season in 2012. Such analysis may add to understanding of the trends associated with the introduction of wolf hunting and trapping seasons, and help improve the stewardship of wolf populations in Minnesota and globally. Statistical analysis showed that wolf mortality was close to constant from 2004 until the first hunting season, followed by a doubling of wolf mortality after the initiation of the first hunting season in 2012, with mortality remaining at a nearly constant elevated level through 2019. The finer-grained trend implies that human-caused mortality increased sharply during hunting seasons, while naturally attributable mortality initially dropped. While gradually diminishing after the discontinuation of hunting, human-caused mortality persisted at a higher level than before the initiation of hunting.

## Results

A regression analysis was applied to MN DNR wolf radiotelemetry data for the 2004–2019 period, as described in the Methods (see Table A1 for a summary of wolf radiotelemetry data by calendar year). The goal was to compare two time periods represented in the data. For brevity, the first period is called “before hunt,” which started in the middle of 2004 when the first two wolves were radio-collared for the MN DNR tracking study, with a larger, more representative number of radio-collared wolves being added between 2005 and October 31st, 2012—before the initiation of the first wolf hunting and trapping season. The second period, which is called “after hunt”, started on November 1st, 2012. The hunting season started in the first week of November, while late hunting and trapping continued in January 2013. Similarly, the second and third annual hunting seasons started in November of each respective year. Thus, statistical comparison years covered from November 1st in one year to October 31st in the next year (see Table A2 for wolf radiotelemetry summary with time reference offset to November 1st). The regression discontinuity analysis, described below, validates the need for this choice of time frames.

Overall, 150 wolves were tracked for a combined total of more than 50,000 radio-days over a period of nearly 16 years. This amount of data resulted in accurate summary survival statistics for the entire period, and separately for before and after hunting periods (see Table 1).

**Table 1:** Wolf survival and mortality summary.

| Period | Survival | Yearly mortality |  |  | R-days | Wolf deaths by cause |  |
| --- | --- | --- | --- | --- | --- | --- | --- |
|  |  | all | human | natural |  | human | natural |
| All wolves |  |  |  |  |  |  |  |
| Before | 0.783 (0.69-0.89) | 21.7% | 10.0% | 11.7% | 22373 | 6.92 | 8.08 |
| After | 0.566 (0.48-0.67) | 43.4% | 35.8% | 7.6% | 28233 | 36.30 | 7.70 |
| All | 0.653 (0.59-0.73) | 34.7% | 25.4% | 9.3% | 50606 | 43.22 | 15.78 |
| Adult |  |  |  |  |  |  |  |
| Before | 0.773 (0.67-0.89) | 22.7% | 11.3% | 11.3% | 18440 | 6.50 | 6.50 |
| After | 0.574 (0.48-0.69) | 42.6% | 33.0% | 9.6% | 22341 | 26.32 | 7.68 |
| All | 0.656 (0.58-0.74) | 34.4% | 24.0% | 10.4% | 40781 | 32.82 | 14.18 |
| Male |  |  |  |  |  |  |  |
| Before | 0.754 (0.61-0.93) | 24.6% | 14.1% | 10.6% | 9035 | 4.00 | 3.00 |
| After | 0.583 (0.45-0.76) | 41.7% | 23.8% | 17.9% | 10841 | 9.14 | 6.86 |
| All | 0.655 (0.55-0.78) | 34.5% | 19.7% | 14.8% | 19876 | 13.14 | 9.86 |
| Female |  |  |  |  |  |  |  |
| Before | 0.792 (0.66-0.95) | 20.8% | 8.3% | 12.5% | 9405 | 2.40 | 3.60 |
| After | 0.565 (0.43-0.74) | 43.5% | 41.0% | 2.6% | 11500 | 16.94 | 1.06 |
| All | 0.658 (0.56-0.78) | 34.2% | 27.6% | 6.6% | 20905 | 19.34 | 4.66 |
| Juvenile |  |  |  |  |  |  |  |
| Before | 0.831 (0.64-1.07) | 16.9% | 0.0% | 16.9% | 3933 | 0.00 | 2.00 |
| After | 0.538 (0.37-0.79) | 46.2% | 46.2% | 0.0% | 5892 | 10.00 | 0.00 |
| All | 0.640 (0.50-0.82) | 36.0% | 30.0% | 6.0% | 9825 | 10.00 | 2.00 |

Wolf hunting seasons always commenced in November of 2012–2014; therefore, all statistics are offset to start on November 1st of each year. The “before” period includes 2004 to October 31, 2012, while the “after” period includes November 1st, 2012 to the end of 2019. All years were calculated for the entire data period, combining before and after periods. The range in parentheses following “Survival” is the 95% confidence interval range. R-days are the number of radiotelemetry days for each time interval. Fractional wolf death counts are the result of six wolf deaths that were of unknown causes, imputed proportionately to the known mortality causes for each period.

As shown in the section titled *All wolves* in Table 1, overall wolf mortality (all wolf ages and sexes) doubled from the before-hunt to after-hunt periods. The trends were similar for the *Adult*, *Male*, and *Female* sections, with male wolves showing slightly less than double the mortality increase from the before to after hunt periods, while overall female wolf mortality more than doubled and juvenile wolf mortality nearly tripled. Human-caused mortality increased for all wolf categories after the first hunting season; however, this increase was most notable for juvenile wolves, followed by female wolves. The proportion of natural cause mortality decreased after the hunt for all categories, with the exception of male wolves, whose natural mortality also increased—potentially due to conflicts caused by wolf pack disruption during and after hunts [13].

Table 1 shows that all-cause mortality across wolf categories was within a close range among demographic groups, with juveniles showing the lowest mortality before the hunt and the highest mortality after the hunt, with all recorded after-hunt mortality being of human causes. Over the entire 16-year span of this study, 23 deaths were recorded for male wolves, 24 for female wolves, and 12 for juveniles. The 95% confidence intervals show the tightest fit for “All wolves” in Table 1. The 95% confidence intervals widened (along with fractionalizing radio-days) when splitting statistics by males, females, and juveniles, therefore discouraging the granular age-based regression on an annual scale separately for these categories. Thus, it made sense to perform the regression analysis on the overall tracked population.

A summary of the linear regression analysis is presented in Fig. 1. The daily unit hazard was calculated for each tracked year, linear regression was performed across the entire period (Fig. 1a), and corresponding yearly survival was derived from this result (Fig. 1b). The survival would be 0.83 in 2004, decreasing to 0.51 in 2019; however, there appears to be a distinct grouping of the survival data points (and daily unit hazard) for the periods before and after the hunt. Regression discontinuity analysis showed a significant trend cutoff point [14] with a p-value of 0.02–0.04 exclusively within the October–December 2012 period (see Regression discontinuity analysis in the Methods and Appendix A.3, particularly Table A9). Independently, the statistical year offset for each 12-month period in Appendix A.2 showed the lowest standard error of linear regression for the statistical year starting on November 1st for all three regression periods: before hunt, after hunt, and even for a single regression over the entire period.

**Fig. 1:**
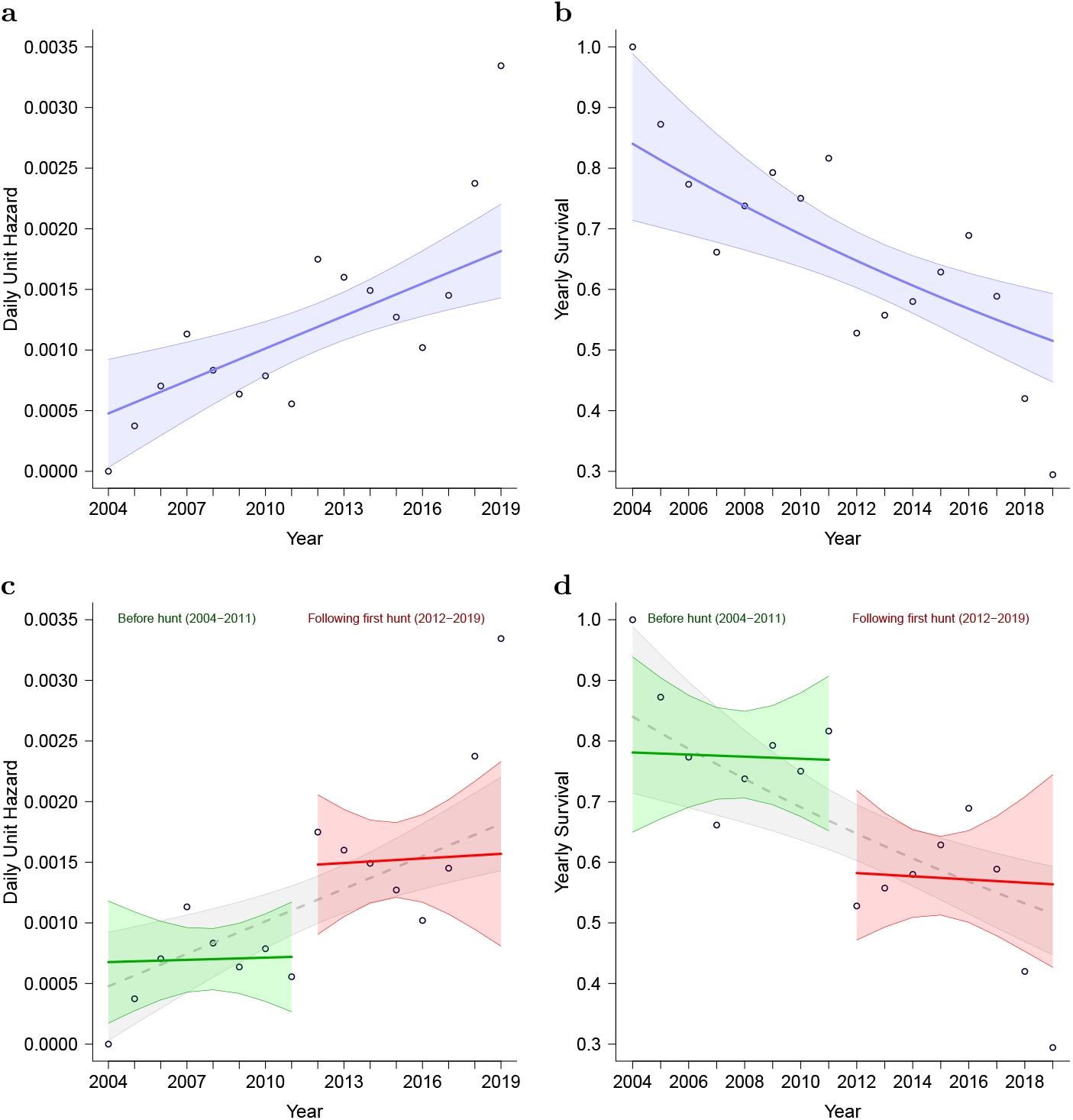
Survival statistics summary. Wolf hunting seasons always commenced in November 2012–2014; therefore, all statistics are offset to start on November 1st of each year. **(a)** Linear regression of daily unit hazard across the entire 2004–2019 period. Blue line – regression; blue shaded area – 95% confidence interval. **(b)** Linear regression of yearly survival across the entire 2004–2019 period. Blue line – regression; blue shaded area – 95% confidence interval. **(c)** Daily unit hazard regressions, separately for before (2004–2011) and after the first hunting season (2012–2019). Green line – trend before hunt, red line – trend after hunt; corresponding shaded areas – 95% confidence interval. **(d)** Yearly survival regressions, separately for before (2004–2011) and after the first hunting season (2012–2019). Green line – trend before hunt, red line – trend after hunt; corresponding shaded areas – 95% confidence interval.

Trends tend to behave differently on both sides of a discontinuity, which called for separate regression analyses for these two periods. The regression analyses in Fig. 1c demonstrated relatively constant hazard/mortality periods before the hunt (shown in green), with daily hazard approximately doubling in the first hunt year and continuing on this new and almost constant hazard level after the hunt (red). A gray outline is retained for comparison with the entire 2004–2019 period regression in Fig. 1a. The corresponding analysis of yearly survival is presented in Fig. 1d, showing two roughly constant survival levels before and after the hunt, with a sharp 22% reduction in survival in 2012.

The mortality analysis presented in Fig. 2a matches the regression discontinuity analysis in Fig. A1b. Fig. 2b presents the analysis with the data points and regression trend separated into human and natural causes, with their corresponding data points and confidence intervals, by cause of death using the overall mortality trend (dashed black line) as a reference. The human-caused and natural mortality numbers were low and close to equal before the hunt, both trending on an approximately constant level. The after-hunt period shows a distinct approximate tripling of human-caused mortality during the first hunt year, which only gradually decreased over the following years, with natural mortality notably dropping in the first hunt year and gradually increasing over the following years, altogether adding up to an approximate doubling of overall wolf mortality over the years following the initiation of hunting and trapping seasons.

**Fig. 2:**
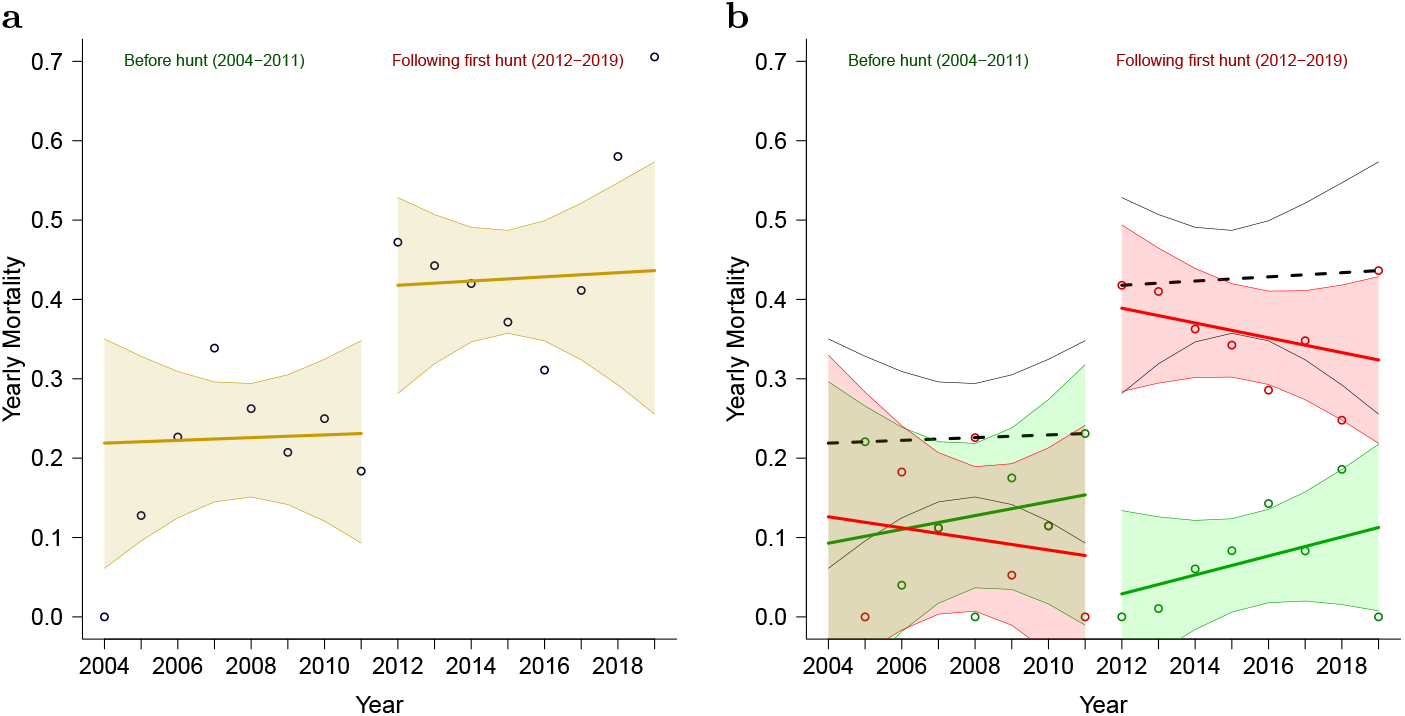
Wolf mortality for the periods before and after the first hunt. **(a)** All-cause mortality before and after hunting seasons. Beige line – trend before and after hunt indicated near the top of the figure; corresponding shaded areas – 95% confidence interval. **(b)** Same as (a), with mortality shared by natural and human causes. Green lines and data points – mortality trend by natural causes, red lines, and data points – mortality trend by human causes; corresponding shaded areas – 95% confidence interval. Black dashed line – trend and confidence interval outlining the sum of all-cause mortality, as shown in plot (a). Before and after hunt periods are indicated near the top of the figure.

## Discussion

In this research, linear regression analysis was applied to ascertain the patterns of wolf survival and mortality in Minnesota before and after the initiation of the 2012–2014 wolf hunting seasons based on wolf radiotelemetry statistics collected by the MN DNR during the 2004–2019 period. The regression discontinuity analysis, regression weighting fit when compared to the high-confidence extended period survival summaries, and the smallest residual standard errors of linear regression achieved when comparing 12-month periods starting on November 1st of each year (see Appendix A.2 and Appendix A.3) all point to statistically significant trend discontinuity timed with the initiation of the first hunting season in November 2012.

The period before wolf hunting seasons was characterized by near-constant wolf mortality averaging at 21.7% (see Table 1, with approximately equal proportions of mortality from human and natural causes (see Fig. 2b). After the initiation of the wolf hunting seasons in 2012, wolf mortality doubled to an average of 43.4% and remained close to constant for the after-hunt period, with human-caused mortality becoming predominant, while the recorded natural mortality precipitously dropped during the wolf hunting seasons of 2012–2014. Human-caused mortality was highest during the hunting seasons, and while it started diminishing later, it remained notably higher than before the initiation of the hunting seasons throughout the year 2019. Notably, although overall mortality increased similarly for all wolf categories, human-caused mortality more than quadrupled for females and juveniles, and their natural mortality nearly vanished initially. For male wolves, while human-caused mortality less than doubled, the natural cause mortality also increased by 70%, which may have been caused by increased conflict and hunting difficulty due to pack disruption [13, 15, 16].

The present study’s findings of a rapid increase in wolf mortality following hunt initiation disagree with the only other recent paper using the same data set that was published by Chakrabarti et al. [12], who “did not observe evidence that survival was markedly reduced during years when a regulated hunting and trapping season was implemented for wolves (years 2012–2014)”. Chakrabarti et al. [12] applied Bayesian analysis in an attempt to determine which of the five smooth regression models tested would best fit the observed data (see Equations (1)–(6) in [12]). Smooth continuous models are not well suited for detecting trend discontinuities, and the model information criteria—DIC, WAIC, and LOOIC [17]—shown in Table 1[12] differed only slightly among these five models, thereby indicating that no one model performed significantly better than any of the others. In Chakrabarti et al.’s [12] defense, as present study showed in Appendix A.3, running a smooth statistical model in 12-month statistical periods based on the calendar year may have been additionally prone to masking discontinuous behavior. We were aided by testing for trend cutoffs and compared regression analysis trends for periods before and after the introduction of the hunts (see Fig. 2). This also allowed us to discover finer changes in the patterns of natural and human-caused wolf mortality.

Human-caused mortality can be due to factors such as hunting, trapping, problem wolf elimination, and roadkill [12, 18]. Unsurprisingly, the level of human-caused mortality increased during hunting and trapping seasons. In 2012 alone, nearly 20% of the estimated wolf population was hunted and trapped [6]. However, mortality remaining high after hunts were discontinued may indicate human attitudes becoming more negative toward wolves after their killing was legalized for a period of time [19–22].

Remarkably, Fuller [23] reported similarly high levels of wolf mortality in a 1980–1985 radio-tracking study of 81 wolves, with even higher human-caused mortality than that observed following the 2012–2014 hunting and trapping seasons—even though the wolf population was smaller in the 1980s. Fuller reported that during times when with “complete federal protection in 1974, controversy over wolves in Minnesota has abounded”[23]. A recent study in Wisconsin and Michigan concluded [19] that liberalizing culling or hunting is more likely to increase illegal killing than reduce it, with controversy abounding again in Minnesota and neighboring states [24–27].

**In conclusion**, before the initiation of the wolf hunting and trapping seasons of 2012–2014, wolf mortality was stable, with yearly wolf mortality being approximately 21.7% and caused by approximately equal proportions by natural and human causes, with natural causes being slightly more common. Something resembling a phase shift occurred with the initiation of the first hunting season, when wolf mortality doubled to 43.4%, became predominantly linked to human causes, and remained on such an elevated level with only a slight gradual decrease over the 5 years following hunt discontinuation. On average, for the period after the initiation of wolf hunting seasons, human causes were linked to 35.8% of the entire wolf population each year, with natural mortality being responsible for 7.6% of the wolf population. This may reflect the reality that with such high yearly human-caused mortality, not many wolves could reach the age when frailty, diseases, and injuries would become the main causes of death, as it appears was the situation before the wolf hunting seasons in 2012–2014.

## Methods

### Description of the radiotelemetry data

MN DNR wolf telemetry data for 2004–2019 was received on request from the authors of Chakrabarti et al. [12] (see Supplementary Data file *MnWolfSurvivalMortalityAge.csv*).

The survival analysis was performed by first aggregating the radio-days for each tracked wolf, with the cause of death date or censoring date parsed and sorted with the help of the C program *WolfDNR.exe*, which was written for this purpose by the author. Similarly to [12], the pups and yearlings were counted together as juveniles for the final statistics since there were only 12 cases of juvenile mortality altogether. The juveniles’ radio-days were accounted for so that April 15th of each year was considered a “graduation day” for the older age category [12, 23]. When crossing April 15th of a year at 2 years of age, these wolves’ radio-days were counted as adult radio-days thereafter. Notably, there was one occasion in the source file when pup W05-2270 was registered on May 6, 2005, at which age it was unlikely to be radio-collared. Thus, it was most likely a yearling, and an adjustment to the program was made to account for such scenarios, although it was the only exceptional case. Therefore, pup W05-2270 was reclassified as a yearling; within less than a year, it was censored while still yearling. The radiotelemetry data summaries parsed and arranged by the C program *WolfDNR.exe* (described further in this section) are visually explanatory, unlike the raw input CSV file. See the summary of a standard calendar year view in Table A1 and statistical year offset to November 1st of each year in Table A2, with corresponding radio-days and mortality counts.

### Imputation of wolf deaths from unknown causes

Of the 59 wolves reported dead in the dataset, the causes of mortality for 6 wolves were not determined. A preliminary analysis showed that mortalities within periods before the hunt and after the hunt remained at a relatively constant level for each of these periods. The ratio of human and natural mortalities was determined for before and after the hunt periods, and the missing mortality cause was imputed by assigning each of these mortalities’ fractional values for human and natural causes. See the comparisons between Table 1 and corresponding Table A11, where unknown deaths were omitted. The patterns appear to be qualitatively the same between the two representations, with 10% lower values across all numbers where the unknown deaths were omitted, which demonstrates the benefit of this imputation (see Appendix A.4).

### Survival analysis

The yearly wolf radio-days were calculated in *WolfDNR.exe* using the equation:

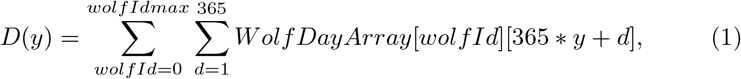

thus summing all entries for each individual wolf ID in the MN DNR data file that were accumulated above in *W olf DayArray* for the continuous count of days *d* of each year starting from 2004 (denoted as *y*), resulting in the sum of *D* for each year. The deaths or censored individuals, with their causes, wolf sexes, and ages, were added for each year in this same loop.

The average daily statistics were calculated using the maximum likelihood estimator (MLE) [28, 29]. The periods were considered tracked daily (see also [12]), and wolf mortality considered established as the end date in the MN DNR data (see further treatment in discussion of regression discontinuity in Appendix A.3). The resulting survival numbers for this simple case scenario matched the initial value of Trent & Rongstad’s [30] daily survival equation in the first MLE iteration:

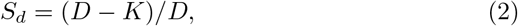

where *D* represents the sum of radio-days for the given period and *K* represents recorded mortality counts during this period.

The yearly survival was counted for standard 365-day years:

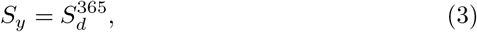

where *S_y_* is yearly survival. Yearly mortality *M_y_* is complementary to yearly survival:

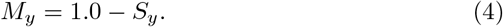

### Weighted linear regression analysis

As the radio-day coverage was not even over the years, additionally statistics recording zero wolf mortality in the statistical year 2004, weighted linear regression was required to prevent bias by less representative data [31]. Two approaches are possible in this data set: weighting by yearly variance or weighting by yearly radio days squared. Both methods resulted in a remarkably close outcome, as discussed in Appendix A.2 (see comparisons in Table A3—Table A8). The weighting by yearly radio-days squared almost precisely matched the higher confidence summary survival data in Table 1, and thus was chosen for this study. See Appendix A.2 for the evaluation and choice of linear regression weighting.

### Regression discontinuity analysis

The R script *WolfProc.R* iterated through each month of five consecutive years (2010–2014) and tested for the existence of regression discontinuity cutoff points [14]. The only sharp trend discontinuity cutoff period was found to span October, November, and December 2012, with significant p-values of 0.022, 0.039, and 0.041, respectively, when the mortality trend jumped to a 22% higher level than before and continued along this post-discontinuity trend line. Thus, the introduction of wolf hunting seasons in November 2012 corresponded with the statistically significant discontinuity in the wolf mortality trend (see in-depth treatment in Appendix A.3).

### Implementation of linear regression analysis

The linear regression with weighting was implemented using standard R libraries with the core regression model as follows:

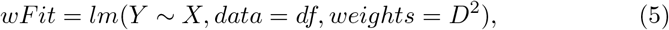

where *wFit* is the weighted fit, Y is the typical daily hazard values, *X* is the year range, *D* is the number of radio-days (measurements) per year, *K* is the recorded number of wolf deaths, and *df* is an R data frame containing input loaded from the CSV data file. A simple linear regression was then used to trend the fraction of mortality by natural and human causes.

### Code and executables

The data preparation and sorting were performed with help of the C program *WolfDNR.exe*, which outputs CSV files with radio-days, mortality counts, corresponding daily hazard (overall and by mortality cause), yearly survival, and mortality data points using the approach described above. The majority of the processing was performed using the R script *WolfProc.R*, which takes as an input the raw yearly summaries aggregated by *WolfDNR.exe*, performs linear regression and regression discontinuity analysis, extracts the relevant statistics, and outputs the figures and tables in PDF and LaTeX formats, as used in the manuscript. Two additional R scripts, *WolfSummaryTableBeforeAfter.R* and *WolfTableByYear.R*, output the remaining tables used in the manuscript. All of the above programs and code are available in the Supplementary Data, along with the batch files that allow to perform all the processing, starting with the MN DNR data set CSV file and then outputting the data tables and graphs used in this manuscript. Although this code is intended for use with the MN DNR data set, it can be easily adjusted for processing different datasets.

## Statistics and reproducibility

The survival analysis was performed on radiotelemetry data originally sourced from MN DNR and obtained on request from Chakrabarti et al. [12]. The linear regression fit and discontinuity analysis were performed using standard R libraries. The reported tables and figures include the 95% confidence intervals. The regression discontinuity analysis used a p-value significance of ≤ 0.05.

## Data availability

The input data, source code, executable, and batch files necessary to perform analysis are available in the Supplementary Data ZIP file.

## Author contributions

R.O. conceived the project, performed the statistical analysis, and wrote the manuscript.

## Appendix A

## A.1 MN DNR radiotelemetry data summary for the 2004–2019 period

**Table A1:** Radiotelemetry summaries for standard calendar years starting **January 1st**.

| Year | R-days | Deaths |  |  | Daily hazard |  |
| --- | --- | --- | --- | --- | --- | --- |
|  |  | all | human | natural | unit | variance |
| 2004 | 469 | 0 | 0.000 | 0.000 | 0e+00 | 0e+00 |
| 2005 | 1668 | 0 | 0.000 | 0.000 | 0e+00 | 0e+00 |
| 2006 | 2946 | 2 | 1.000 | 1.000 | 6.79e-04 | 2.3e-07 |
| 2007 | 4645 | 4 | 3.545 | 0.455 | 8.61e-04 | 1.85e-07 |
| 2008 | 2716 | 3 | 1.000 | 2.000 | 1.1e-03 | 4.06e-07 |
| 2009 | 1839 | 0 | 0.000 | 0.000 | 0e+00 | 0e+00 |
| 2010 | 3030 | 2 | 0.545 | 1.455 | 6.6e-04 | 2.18e-07 |
| 2011 | 2175 | 2 | 1.000 | 1.000 | 9.2e-04 | 4.22e-07 |
| 2012 | 4412 | 5 | 3.000 | 2.000 | 1.13e-03 | 2.57e-07 |
| 2013 | 2889 | 6 | 5.786 | 0.214 | 2.08e-03 | 7.17e-07 |
| 2014 | 4617 | 6 | 6.000 | 0.000 | 1.3e-03 | 2.81e-07 |
| 2015 | 4424 | 5 | 4.000 | 1.000 | 1.13e-03 | 2.55e-07 |
| 2016 | 4487 | 8 | 5.786 | 2.214 | 1.78e-03 | 3.97e-07 |
| 2017 | 3316 | 2 | 2.000 | 0.000 | 6.03e-04 | 1.82e-07 |
| 2018 | 4716 | 9 | 6.571 | 2.429 | 1.91e-03 | 4.04e-07 |
| 2019 | 2388 | 5 | 3.000 | 2.000 | 2.09e-03 | 8.75e-07 |

The R-days column title stands for radiotelemetry days. Fractional wolf death counts are the result of six wolf deaths with unknown causes, imputed proportionately to the known causes of mortality for each period.

**Table A2:** Radiotelemetry summaries with the first day of the year offset to **November 1st**, the month of wolf hunting season commencement.

| Year | R-days | Deaths |  |  | Daily hazard |  |
| --- | --- | --- | --- | --- | --- | --- |
|  |  | all | human | natural | unit | variance |
| 2004 | 1424 | 0 | 0.000 | 0.000 | 0e+00 | 0e+00 |
| 2005 | 2672 | 1 | 0.000 | 1.000 | 3.74e-04 | 1.4e-07 |
| 2006 | 4262 | 3 | 2.462 | 0.538 | 7.04e-04 | 1.65e-07 |
| 2007 | 3533 | 4 | 2.000 | 2.000 | 1.13e-03 | 3.2e-07 |
| 2008 | 1200 | 1 | 1.000 | 0.000 | 8.33e-04 | 6.94e-07 |
| 2009 | 3143 | 2 | 0.462 | 1.538 | 6.36e-04 | 2.02e-07 |
| 2010 | 2541 | 2 | 1.000 | 1.000 | 7.87e-04 | 3.1e-07 |
| 2011 | 3598 | 2 | 0.000 | 2.000 | 5.56e-04 | 1.54e-07 |
| 2012 | 3431 | 6 | 6.000 | 0.000 | 1.75e-03 | 5.09e-07 |
| 2013 | 4375 | 7 | 6.825 | 0.175 | 1.6e-03 | 3.65e-07 |
| 2014 | 4694 | 7 | 6.000 | 1.000 | 1.49e-03 | 3.17e-07 |
| 2015 | 4720 | 6 | 4.825 | 1.175 | 1.27e-03 | 2.69e-07 |
| 2016 | 2941 | 3 | 2.000 | 1.000 | 1.02e-03 | 3.46e-07 |
| 2017 | 4825 | 7 | 5.650 | 1.350 | 1.45e-03 | 3e-07 |
| 2018 | 2948 | 7 | 4.000 | 3.000 | 2.37e-03 | 8.04e-07 |
| 2019 | 299 | 1 | 1.000 | 0.000 | 3.34e-03 | 1.11e-05 |

Wolf hunting seasons always commenced in November of 2012–2014; therefore, all statistics are offset to start each year on November 1st. This data arrangement was used as the basis for the regression analysis in this study. The R-days column title stands for radiotelemetry days. Fractional wolf death counts are the result of six wolf deaths with unknown causes, imputed proportionately to the known causes of mortality for each period.

## A.2 Fit of weighting methods used in linear regression

The MN DNR radiotelemetry data were characterized by years with varying numbers of cumulative radio-days and wolf mortality events (as shown in Table A1 and Table A2). A common approach to limit bias from less representative data samples and outliers involves applying linear regression with weighting [31]. In statistical sampling, the inverse of variance is often used. However, in our data, years 2004, 2005, and 2009 in Table A1 have 0 wolf deaths. Thus, no variance was calculated. Similarly, the year 2004 in Table A2 has 0 wolf deaths (no variance) and quite a low number of radio-days, with one wolf death producing an outlier in the year 2019. Although the outliers were handled well by the variance weighting, the zero variance required an additional estimation of possible variance, perhaps from nearby valid entries. Additionally, with high-granularity radiotelemetry (as used in MN DNR data collection), the greater variance in years with higher wolf mortality may create its own bias. A second weighting approach involves using radio-days:

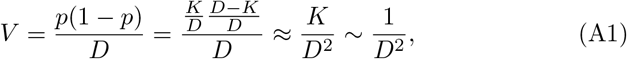

where *V* is variance, *p* and (1 − *p*) are the probability fractions, *D* is the number of radio-days (measurements), and *K* is the count of wolf deaths in the radio-days period. With a large *D* and small *K*, (*D − K*)/*D* ≈ 1, and *K/D*^2^ ∼ 1/*D*^2^. The final step shows the use of the unbiased number of recorded deaths weighted by radio-days; thus, when weighting can be performed as 1/*V*, it must be performed alternatively as *D*^2^. The advantage of this second weighting approach is its consistency, and it does not require inventing a special treatment for the years with no recorded wolf mortality. The following comparisons show that this weighting method also allows a more precise match of the regression to the summary analysis in Table 1.

Finally, the R models for these two weights are implemented as follows:

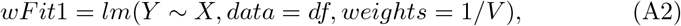

and

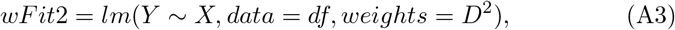

where wFit1 and wFit2 are the corresponding weighted fit scenarios for the linear regression, Y are the typical daily hazard values or survival, X is the year range, D is the number of radio-days (measurements) per year, and df is an R data frame with a loaded CSV input (see also Sheather [31]).

On the following three pages, the tables are grouped in comparison pairs for weighting using inverse variance and *D*^2^ (see Table A3–Table A4, Table A5–Table A6, and Table A7–Table A8). For completeness, these tables show the parameters of the linear regression with years offset to the first day of each month of the year. The most important columns are stdError (the standard error of the regression), *r*^2^ (the goodness-of-fit), and mean survival, which is typically calculated as:

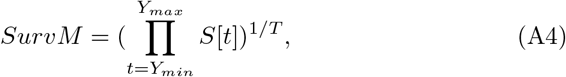

where *SurvM* is the mean survival over the subsequent number of years T in the regression interval *Y_min_* to *Y_max_*. As shown below, stdErr is comparatively lowest for years offset to start as of November 1st in all tables except Table A4, where it is a close second. The regression parameter *r*^2^ is a goodness-of-fit of the trend. In Table A3–Table A6 *r*^2^ is low, representing a near-constant level of survival and mortality. For the entire dataset duration the best regression was also observed with a November 1st time frame offset.

The second equally important consideration for the weighting method is how well the mean survivals in these tables match the multiyear summaries in Table 1. Here, the summaries from sections in Table 1 compared to radio-days squared weighting in the regression analysis, are presented as follows (Equation A3 scenario): for the 2004–2011 period, 0.783 vs 0.775; 2012–2019, 0.566 vs 0.573; all years’ time intervals, 0.653 vs 0.658 — almost exact matches. Inverse variance weighting (Equation A2 scenario) diverged further from the summary values in Table 1—showing 0.783 vs 0.804, 0.566 vs 0.582, and 0.653 vs 0.685, respectively—while exaggerating the summary survival in each case by a few percent.

Consequently, the regression analysis when using inverse variance weighting resulted in nearly identical but slightly underestimated mortality trend values. Thus, we used the weighting by radio-days squared in our reported analysis.

**Table A3:**
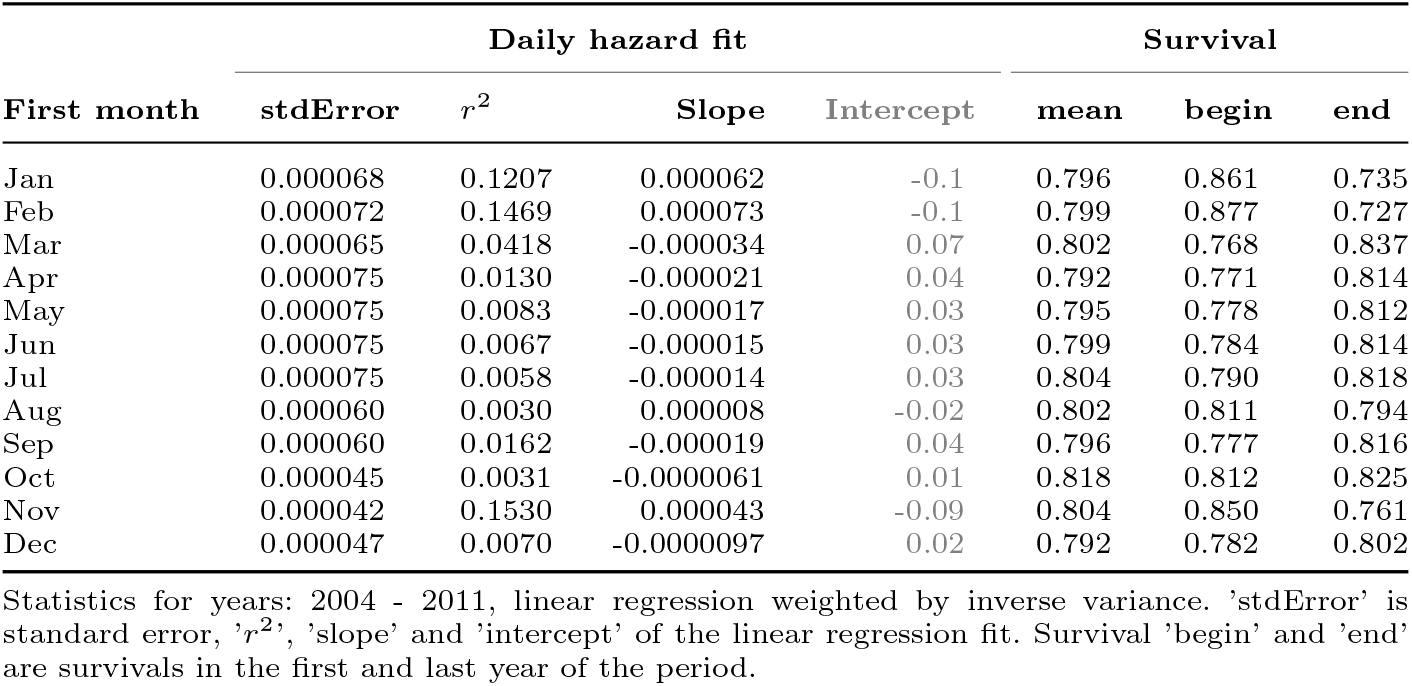
Linear regression fit parameters by assigned first month of the year for **years 2004–2011, inverse variance weighting**.

**Table A4:**
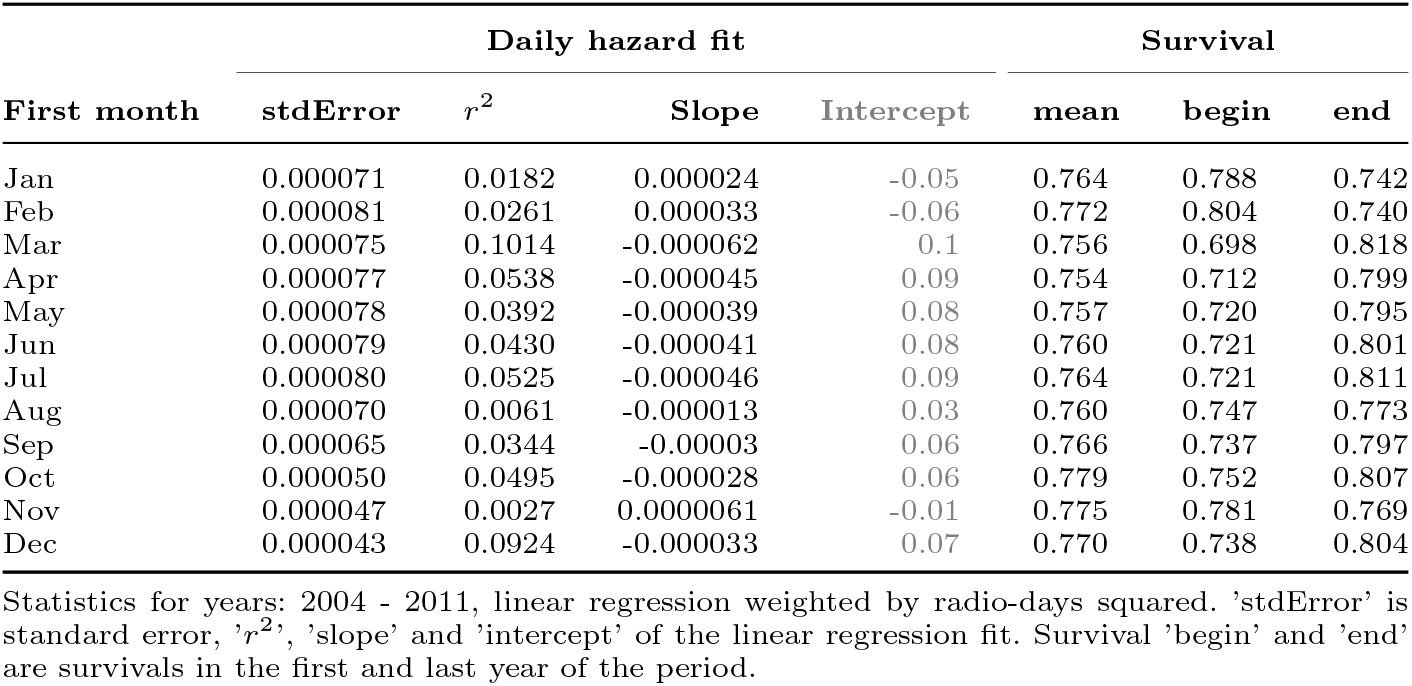
Linear regression fit parameters by assigned first month of the year for **years 2004–2011, radio-days squared weighting**.

| First month | Daily hazard fit |  |  |  | Survival |  |  |
| --- | --- | --- | --- | --- | --- | --- | --- |
| | stdError | $r^2$ | Slope | Intercept | mean | begin | end |
| Jan | 0.000071 | 0.0182 | 0.000024 | -0.05 | 0.764 | 0.788 | 0.742 |
| Feb | 0.000081 | 0.0261 | 0.000033 | -0.06 | 0.772 | 0.804 | 0.740 |
| Mar | 0.000075 | 0.1014 | -0.000062 | 0.1 | 0.756 | 0.698 | 0.818 |
| Apr | 0.000077 | 0.0538 | -0.000045 | 0.09 | 0.754 | 0.712 | 0.799 |
| May | 0.000078 | 0.0392 | -0.000039 | 0.08 | 0.757 | 0.720 | 0.795 |
| Jun | 0.000079 | 0.0430 | -0.000041 | 0.08 | 0.760 | 0.721 | 0.801 |
| Jul | 0.000080 | 0.0525 | -0.000046 | 0.09 | 0.764 | 0.721 | 0.811 |
| Aug | 0.000070 | 0.0061 | -0.000013 | 0.03 | 0.760 | 0.747 | 0.773 |
| Sep | 0.000065 | 0.0344 | -0.00003 | 0.06 | 0.766 | 0.737 | 0.797 |
| Oct | 0.000050 | 0.0495 | -0.000028 | 0.06 | 0.779 | 0.752 | 0.807 |
| Nov | 0.000047 | 0.0027 | 0.0000061 | -0.01 | 0.775 | 0.781 | 0.769 |
| Dec | 0.000043 | 0.0924 | -0.000033 | 0.07 | 0.770 | 0.738 | 0.804 |

Statistics for years: 2004 - 2011, linear regression weighted by radio-days squared. 'stdError' is standard error, ' $r^2$ ', 'slope' and 'intercept' of the linear regression fit. Survival 'begin' and 'end' are survivals in the first and last year of the period.

**Table A5:** Linear regression fit parameters by assigned first month of the year for **years 2012–2019, inverse variance weighting**.

| First month | Daily hazard fit |  |  |  | Survival |  |  |
| --- | --- | --- | --- | --- | --- | --- | --- |
| | stdError | $r^2$ | Slope | Intercept | mean | begin | end |
| Jan | 0.000096 | 0.0041 | 0.000015 | -0.03 | 0.625 | 0.638 | 0.613 |
| Feb | 0.000113 | 0.0069 | -0.000023 | 0.05 | 0.638 | 0.620 | 0.658 |
| Mar | 0.000139 | 0.0101 | 0.000034 | -0.07 | 0.657 | 0.686 | 0.629 |
| Apr | 0.000104 | 0.0019 | 0.000011 | -0.02 | 0.625 | 0.634 | 0.616 |
| May | 0.000109 | 0.0018 | 0.000011 | -0.02 | 0.628 | 0.637 | 0.619 |
| Jun | 0.000113 | 0.0012 | -0.0000098 | 0.02 | 0.626 | 0.618 | 0.634 |
| Jul | 0.000090 | 0.0005 | -0.0000049 | 0.01 | 0.601 | 0.597 | 0.604 |
| Aug | 0.000088 | 0.0054 | 0.000016 | -0.03 | 0.601 | 0.614 | 0.589 |
| Sep | 0.000092 | 0.0028 | 0.000012 | -0.02 | 0.597 | 0.606 | 0.588 |
| Oct | 0.000087 | 0.0007 | -0.0000058 | 0.01 | 0.587 | 0.583 | 0.592 |
| Nov | 0.000079 | 0.0003 | 0.0000034 | -0.005 | 0.582 | 0.585 | 0.580 |
| Dec | 0.000119 | 0.0017 | -0.000012 | 0.03 | 0.593 | 0.584 | 0.603 |

Statistics for years: 2012 - 2019, linear regression weighted by inverse variance. 'stdError' is standard error, ' $r^2$ ', 'slope' and 'intercept' of the linear regression fit. Survival 'begin' and 'end' are survivals in the first and last year of the period.

**Table A6:** Linear regression fit parameters by assigned first month of the year for **years 2012–2019, radio-days squared weighting**.

| First month | Daily hazard fit |  |  |  | Survival |  |  |
| --- | --- | --- | --- | --- | --- | --- | --- |
| | stdError | $r^2$ | Slope | Intercept | mean | begin | end |
| Jan | 0.000079 | 0.1374 | 0.000077 | -0.2 | 0.584 | 0.644 | 0.529 |
| Feb | 0.000098 | 0.0330 | 0.000044 | -0.09 | 0.574 | 0.608 | 0.543 |
| Mar | 0.000107 | 0.0038 | 0.000016 | -0.03 | 0.581 | 0.593 | 0.569 |
| Apr | 0.000097 | 0.0171 | 0.000031 | -0.06 | 0.573 | 0.597 | 0.551 |
| May | 0.000106 | 0.0092 | 0.000025 | -0.05 | 0.573 | 0.592 | 0.555 |
| Jun | 0.000106 | 0.0054 | 0.000019 | -0.04 | 0.582 | 0.596 | 0.568 |
| Jul | 0.000088 | 0.0010 | 0.0000067 | -0.01 | 0.582 | 0.587 | 0.577 |
| Aug | 0.000084 | 0.0092 | 0.00002 | -0.04 | 0.587 | 0.602 | 0.573 |
| Sep | 0.000084 | 0.0003 | 0.0000038 | -0.006 | 0.584 | 0.587 | 0.581 |
| Oct | 0.000080 | 0.0081 | -0.000018 | 0.04 | 0.584 | 0.571 | 0.597 |
| Nov | 0.000069 | 0.0055 | 0.000013 | -0.02 | 0.573 | 0.582 | 0.564 |
| Dec | 0.000117 | 0.0040 | -0.000018 | 0.04 | 0.569 | 0.556 | 0.583 |

Statistics for years: 2012 - 2019, linear regression weighted by radio-days squared. 'stdError' is standard error, ' $r^2$ ', 'slope' and 'intercept' of the linear regression fit. Survival 'begin' and 'end' are survivals in the first and last year of the period.

**Table A7:** Linear regression fit parameters by assigned first month of the year for **years 2004–2019, inverse variance weighting**

| First month | Daily hazard fit |  |  |  | Survival |  |  |
| --- | --- | --- | --- | --- | --- | --- | --- |
| | stdError | $r^2$ | Slope | Intercept | mean | begin | end |
| Jan | 0.000027 | 0.3511 | 0.000074 | -0.1 | 0.704 | 0.862 | 0.575 |
| Feb | 0.000030 | 0.2679 | 0.000067 | -0.1 | 0.712 | 0.856 | 0.593 |
| Mar | 0.000035 | 0.1838 | 0.000063 | -0.1 | 0.737 | 0.874 | 0.621 |
| Apr | 0.000031 | 0.2591 | 0.000069 | -0.1 | 0.707 | 0.853 | 0.586 |
| May | 0.000032 | 0.2480 | 0.000068 | -0.1 | 0.708 | 0.854 | 0.587 |
| Jun | 0.000032 | 0.2432 | 0.000069 | -0.1 | 0.709 | 0.855 | 0.587 |
| Jul | 0.000030 | 0.3481 | 0.000082 | -0.2 | 0.698 | 0.873 | 0.557 |
| Aug | 0.000028 | 0.3981 | 0.000084 | -0.2 | 0.701 | 0.883 | 0.556 |
| Sep | 0.000030 | 0.3567 | 0.000084 | -0.2 | 0.702 | 0.884 | 0.558 |
| Oct | 0.000027 | 0.4305 | 0.000087 | -0.2 | 0.705 | 0.896 | 0.555 |
| Nov | 0.000021 | 0.5959 | 0.000095 | -0.2 | 0.685 | 0.889 | 0.528 |
| Dec | 0.000030 | 0.3397 | 0.000079 | -0.2 | 0.693 | 0.861 | 0.558 |

Statistics for years: 2004 - 2019, linear regression weighted by inverse variance. 'stdError' is standard error, ' $r^2$ ', 'slope' and 'intercept' of the linear regression fit. Survival 'begin' and 'end' are survivals in the first and last year of the period. .

**Table A8:** Linear regression fit parameters by assigned first month of the year for **years 2012–2019, radio-days squared weighting**.

| First month | Daily hazard fit |  |  |  | Survival |  |  |
| --- | --- | --- | --- | --- | --- | --- | --- |
| | stdError | $r^2$ | Slope | Intercept | mean | begin | end |
| Jan | 0.000026 | 0.4570 | 0.000088 | -0.2 | 0.667 | 0.849 | 0.525 |
| Feb | 0.000032 | 0.3626 | 0.000089 | -0.2 | 0.660 | 0.843 | 0.517 |
| Mar | 0.000035 | 0.2356 | 0.000073 | -0.1 | 0.654 | 0.798 | 0.536 |
| Apr | 0.000032 | 0.2998 | 0.000079 | -0.2 | 0.649 | 0.805 | 0.523 |
| May | 0.000035 | 0.2722 | 0.000079 | -0.2 | 0.649 | 0.806 | 0.522 |
| Jun | 0.000034 | 0.2565 | 0.000076 | -0.2 | 0.655 | 0.806 | 0.532 |
| Jul | 0.000031 | 0.3060 | 0.000076 | -0.2 | 0.656 | 0.808 | 0.532 |
| Aug | 0.000028 | 0.3471 | 0.000076 | -0.2 | 0.660 | 0.811 | 0.536 |
| Sep | 0.000028 | 0.3526 | 0.000078 | -0.2 | 0.659 | 0.815 | 0.533 |
| Oct | 0.000027 | 0.3907 | 0.000081 | -0.2 | 0.662 | 0.826 | 0.530 |
| Nov | 0.000023 | 0.5217 | 0.000089 | -0.2 | 0.658 | 0.840 | 0.515 |
| Dec | 0.000033 | 0.3148 | 0.000084 | -0.2 | 0.652 | 0.821 | 0.518 |

Statistics for years: 2004 - 2019, linear regression weighted by radio-days squared. 'stdError' is standard error, ' $r^2$ ', 'slope' and 'intercept' of the linear regression fit. Survival 'begin' and 'end' are survivals in the first and last year of the period.

## A.3 Pinpointing the trend change timing via regression discontinuity analysis

Regression discontinuity analysis was performed to determine whether there was a cutoff point where the trend may have become discontinuous [14], as could occur before and after a specific date or event, with trends having the potential to significantly change in their magnitude and slope. This was performed using the R script *WolfProc.R* as a varying slope regression discontinuity analysis [32–34].

**Table A9:** Regression discontinuity analysis found significant trend cutoff in October, November, and December 2012.

| Cut-point<br>Month | Year |  |  |  |  |
| --- | --- | --- | --- | --- | --- |
|  | 2010 | 2011 | 2012 | 2013 | 2014 |
| Jan | 0.990 (0%) | 0.530 (9%) | 0.520 (9%) | 0.652 (5%) | 0.481 (-7%) |
| Feb | 0.736 (8%) | 0.414 (15%) | 0.470 (13%) | 0.350 (-12%) | 0.443 (-10%) |
| Mar | 0.822 (-5%) | 0.346 (19%) | 0.240 (23%) | 0.656 (-7%) | 0.731 (-5%) |
| Apr | 0.861 (-4%) | 0.646 (8%) | 0.223 (21%) | 0.736 (-4%) | 0.827 (-3%) |
| May | 0.855 (-4%) | 0.671 (8%) | 0.241 (21%) | 0.828 (-3%) | 0.955 (-1%) |
| Jun | 0.825 (-5%) | 0.728 (6%) | 0.205 (22%) | 0.751 (-4%) | 0.426 (-10%) |
| Jul | 0.774 (-5%) | 0.746 (5%) | 0.105 (24%) | 0.766 (-4%) | 0.257 (-13%) |
| Aug | 0.731 (-6%) | 0.745 (5%) | 0.193 (17%) | 0.834 (-2%) | 0.284 (-11%) |
| Sep | 0.971 (-1%) | 0.637 (7%) | 0.095 (21%) | 1.000 (0%) | 0.523 (-7%) |
| Oct | 0.999 (0%) | 0.684 (6%) | 0.022* (25%) | 0.834 (2%) | 0.356 (-9%) |
| Nov | 0.907 (-2%) | 0.813 (3%) | 0.039* (19%) | 0.684 (4%) | 0.460 (-6%) |
| Dec | 0.790 (4%) | 0.721 (6%) | 0.041* (24%) | 0.757 (-4%) | 0.511 (-7%) |

R script tested for the existence of discontinuity cutoffs in each month of five consecutive years. The table cells show the p-value significance of discontinuity, followed in parentheses by the overall wolf mortality percentage change at the discontinuity time. \* marks p-values $\leq 0.05$ .

The R script iterated through each month of five consecutive years and tested for the existence of discontinuity cutoff points, see Table A9. The only sharp trend discontinuity cutoff [14] was found spanning October, November, and December 2012, with significant p-values of 0.022, 0.039, and 0.041, respectively, when the mortality trend jumped to a new 22% higher level (an average of the aforementioned 3 months). This timing coincided with the initiation of the first wolf hunting and trapping season of 2012. There are no candidates for any other timing of cutoffs since the p-values became increasingly less significant for the earlier months of 2012, while all columns for years 2010, 2011, 2013, and 2014 have large p-values.

The January row in Table A9 shows that statistics aligned with standard calendar years starting on January 1st were particularly unsuitable for noticing sharp discontinuity. Only part of the mortality from the 2012 to 2013 hunting season occurred in November and December 2012, thus producing a smooth transition when counting statistics in calendar years.

Fig. A1 graphically illustrates the numerical p-values in Table A9. Testing for the cutoff points on November 1st of years 2011 and 2013 showed a low likelihood of such cut point, with p-values of 0.813 and 0.684, respectively, leading to the appearance of Fig. A1a and Fig. A1c. On the other hand, the p-value was 0.039 for November 2012 and corresponds to a sharp breaking of the trend in Fig. A1b.

**Fig. A1:**
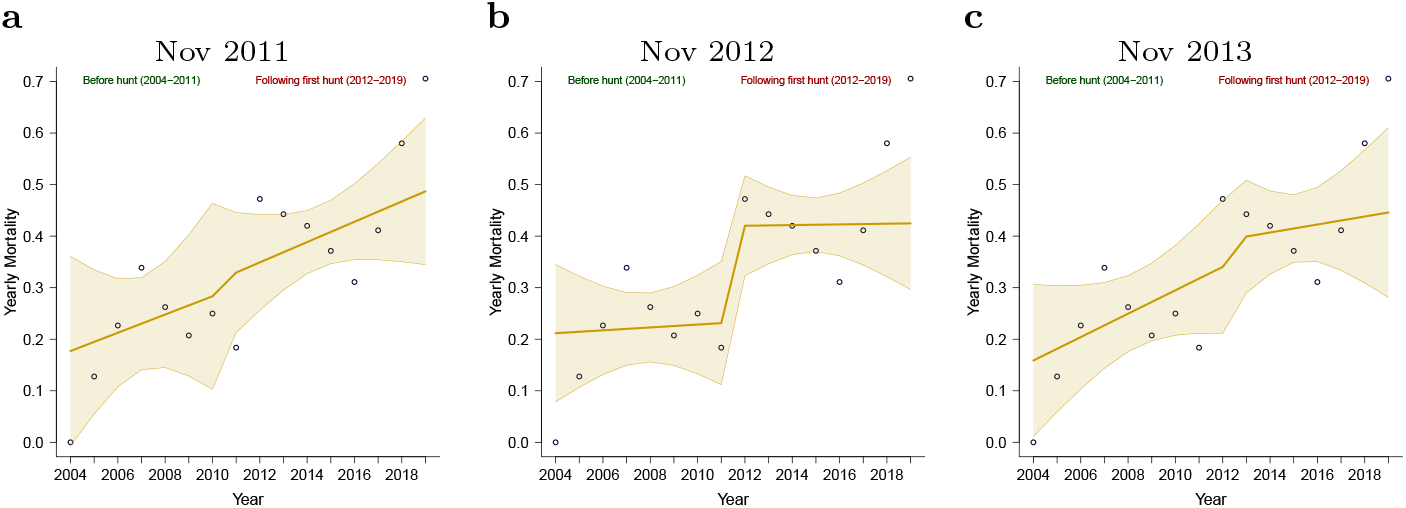
Graphical illustration of the regression discontinuity test results. **(a)** Trend cutoff November 1st, 2011. **(b)** Trend cutoff November 1st, 2012. **(c)** Trend cutoff November 1st, 2013. With the statistical beginning of the year offset to November 1st, large p-values were produced by discontinuity tests for 2011 and 2013, as reflected in the lack of discontinuity in plots (a) and (c). Plot (b) corresponds to a significantly low p-value (0.039) produced by the test for 2012, the year of hunting initiation (see all values in Table A9). The solid beige line shows the mean value of the discontinuous regression, while shaded areas show 95% confidence intervals.

Thus, November 1st, 2012, being the initiation point for the 2012–2014 wolf hunting seasons, coincides well with the middle of the trend discontinuity cutoff in October–December 2012. A choice of October 2012, with the lowest discontinuity p-value (0.022), would result in a similar trend, with an even higher jump in mortality at the trend cutoff point (see Fig. A2).

**Fig. A2:**
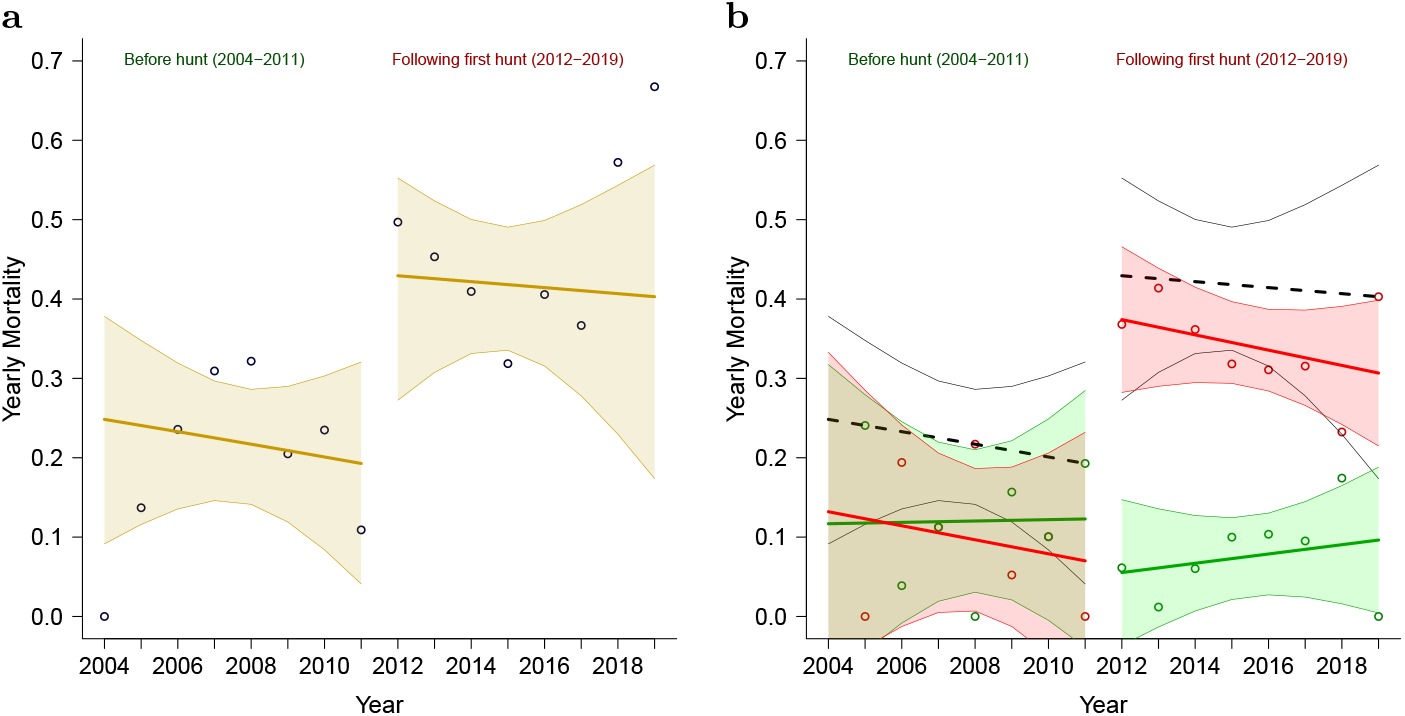
Wolf mortality for the periods before and after the first hunt, using October 1st, 2012 as the trend cutoff point. **(a)** All-cause mortality before and after hunting seasons. Beige line – trend before and after hunting seasons indicated near the top of the figure. **(b)** Same as (a), with mortality shared by natural and human causes. Green lines and data points – mortality trend by natural causes (red lines) and mortality trend by human causes (data points). Black dashed line – trend and confidence interval outlines the sum of all-cause mortality. All corresponding shaded areas – 95% confidence intervals. For comparison with Fig. 2.

There is always an uncertainty inherent to radiotelemetry data when accounting for the fate of the censored wolves and the exact dating of the deaths that may lead to mismeasured wolf mortality [35]. Our analysis used the end date in the MN DNR data for the established wolf death date. There may be other choices of adjusting the uncertainty of the death date, as was done by Chakrabarti et al. [12], however we found that for this data set such adjustments have minimal effect on the survival/mortality results, and any adjustment method without the precise timing knowledge remains arbitrary. A validation using the last seen alive date as the mortality date was performed, demonstrating that even with such extreme edge case adjustment the overall mortality would change by only a fraction of a percent (from 34.7% to 35.6%), and confirming a similar trend discontinuity in October–November 2012 (see in Table A10). Thus, the end date in MN DNR data set was chosen for consistency of treatment with the censored wolves.

**Table A10:** Regression discontinuity validation in edge case of using the last seen dates as the dates of death also found significant trend cutoff in October and November 2012.

| Cut-point<br>Month | Year |  |  |  |  |
| --- | --- | --- | --- | --- | --- |
|  | 2010 | 2011 | 2012 | 2013 | 2014 |
| Jan | 0.912 (2%) | 0.409 (13%) | 0.723 (5%) | 0.550 (-7%) | 0.437 (-9%) |
| Feb | 0.644 (9%) | 0.710 (6%) | 0.756 (5%) | 0.317 (-12%) | 0.458 (-8%) |
| Mar | 0.971 (-1%) | 0.258 (22%) | 0.342 (19%) | 0.750 (-5%) | 0.841 (-3%) |
| Apr | 0.964 (1%) | 0.519 (11%) | 0.391 (15%) | 0.791 (-3%) | 0.903 (-1%) |
| May | 0.971 (1%) | 0.520 (11%) | 0.393 (15%) | 0.880 (-2%) | 0.710 (-4%) |
| Jun | 0.931 (1%) | 0.357 (11%) | 0.165 (17%) | 0.999 (0%) | 0.523 (-6%) |
| Jul | 0.978 (0%) | 0.437 (10%) | 0.160 (17%) | 0.865 (-2%) | 0.232 (-11%) |
| Aug | 0.937 (1%) | 0.303 (11%) | 0.205 (13%) | 0.766 (2%) | 0.462 (-5%) |
| Sep | 0.927 (1%) | 0.364 (10%) | 0.174 (13%) | 0.908 (1%) | 0.546 (-5%) |
| Oct | 0.623 (6%) | 0.284 (11%) | 0.046* (17%) | 0.833 (2%) | 0.230 (-8%) |
| Nov | 0.314 (10%) | 0.665 (4%) | 0.026* (17%) | 0.346 (7%) | 0.630 (-3%) |
| Dec | 0.503 (9%) | 0.835 (3%) | 0.277 (11%) | 0.459 (-7%) | 0.336 (-8%) |

R script tested for the existence of discontinuity cutoff points in each month of five consecutive years. The table cells show the p-value significance of discontinuity, followed in parentheses by the overall wolf mortality percentage change at the discontinuity time. \* marks p-values $\leq 0.05$ .

## A.4 Confirming the consistency of regression when wolf deaths by unknown causes are excluded

Table A11 presents the regression analysis performed while discounting 6 out of 59 wolf mortality events due to unknown causes, rather than imputing them as fractional shares of mortalities. Imputing the unknown cause mortality allows us to account for all wolf deaths in the data rather than underestimating wolf mortality, with no noticeable bias in the regression outcome.

See the comparisons between Table A11 and corresponding Table 1 (where imputation was performed), which show that the patterns appear qualitatively the same between the two representations, with 10% lower values across all numbers where the unknown deaths were omitted, thereby showing the benefit of this imputation.

**Table A11:** Wolf survival and mortality summary **excluding** the imputation of wolf deaths with unknown causes.

| Period | Survival | Yearly mortality |  |  | R-days | Wolf deaths by cause |  |
| --- | --- | --- | --- | --- | --- | --- | --- |
|  |  | all | human | natural |  | human | natural |
| All wolves |  |  |  |  |  |  |  |
| Before | 0.809 (0.72-0.91) | 19.1% | 8.8% | 10.3% | 22373 | 6.00 | 7.00 |
| After | 0.596 (0.51-0.70) | 40.4% | 33.3% | 7.1% | 28233 | 33.00 | 7.00 |
| All | 0.682 (0.62-0.76) | 31.8% | 23.4% | 8.4% | 50606 | 39.00 | 14.00 |
| Adult |  |  |  |  |  |  |  |
| Before | 0.789 (0.69-0.90) | 21.1% | 10.6% | 10.6% | 18440 | 6.00 | 6.00 |
| After | 0.602 (0.50-0.72) | 39.8% | 30.8% | 9.0% | 22341 | 24.00 | 7.00 |
| All | 0.680 (0.61-0.76) | 32.0% | 22.3% | 9.7% | 40781 | 30.00 | 13.00 |
| Male |  |  |  |  |  |  |  |
| Before | 0.754 (0.61-0.93) | 24.6% | 14.1% | 10.6% | 9035 | 4.00 | 3.00 |
| After | 0.624 (0.49-0.80) | 37.6% | 21.5% | 16.1% | 10841 | 8.00 | 6.00 |
| All | 0.680 (0.58-0.80) | 32.0% | 18.3% | 13.7% | 19876 | 12.00 | 9.00 |
| Female |  |  |  |  |  |  |  |
| Before | 0.824 (0.69-0.98) | 17.6% | 7.1% | 10.6% | 9405 | 2.00 | 3.00 |
| After | 0.583 (0.45-0.75) | 41.7% | 39.3% | 2.5% | 11500 | 16.00 | 1.00 |
| All | 0.681 (0.58-0.80) | 31.9% | 26.1% | 5.8% | 20905 | 18.00 | 4.00 |
| Juvenile |  |  |  |  |  |  |  |
| Before | 0.911 (0.76-1.09) | 8.9% | 0.0% | 8.9% | 3933 | 0.00 | 1.00 |
| After | 0.572 (0.40-0.82) | 42.8% | 42.8% | 0.0% | 5892 | 9.00 | 0.00 |
| All | 0.690 (0.55-0.87) | 31.0% | 27.9% | 3.1% | 9825 | 9.00 | 1.00 |

Wolf hunting seasons always commenced in November of wolf hunting seasons 2012–2014; therefore, all statistics are offset to start each year on November 1st. Periods before hunting include 2004 to October 31, 2012, while periods after hunting commenced include November 1st, 2012 to the end of 2019. All years were calculated for the entire data period, combining before and after periods. The range in braces following ‘Survival’ is a 95% confidence interval range. R-days stands for the number of radiotelemetry days for each time interval. Fractional wolf death counts are the result of six wolf deaths with unknown causes, imputed proportionately to the known mortality causes for each period.

