## Supplementary material for "Human-caused wolf mortality persists for years after discontinuation of hunting": Header.h

```
#pragma once


class CLocalMemoryObject
{
public:
	void * m_pMemory;
	__int64 m_iMemorySize;
	CLocalMemoryObject( DWORD dAllocationType, __int64 iMemorySize )
	{
		m_iMemorySize = iMemorySize;
		if( iMemorySize ) {
			m_pMemory = GlobalAllocPtr( dAllocationType, m_iMemorySize );
		} else {
			m_pMemory = NULL;
		}
	};
	~CLocalMemoryObject() 
	{
		if( m_pMemory ) {
			GlobalFreePtr( m_pMemory );
			m_pMemory = NULL;
		}
	};
};
```
