## Supplementary material for "Human-caused wolf mortality persists for years after discontinuation of hunting": PROCESS and BUILD ALL Tables and Figures.bat

```
call RunExe.bat NoPause

call RunSummaries.bat NoPause

cd .\R

call ProcessR.bat NoPause

call ProcessR-Tables.bat NoPause

call ProcessR-YearlyTables.bat NoPause

REM IF All figures and tables are prepared without errors, RUN BUILD of WolvesBeforeAfterHunt.tex in .\LATEX folder

PAUSE
```
