## Supplementary material for "Human-caused wolf mortality persists for years after discontinuation of hunting": ProcessR-Tables.bat

```
rscript.exe WolfSummaryTableBeforeAfter.R "./summaries" "../LATEX/Tables"  "Nov"

rscript.exe WolfSummaryTableBeforeAfter.R "./summaries" "../LATEX/Tables"  "NNov"

rscript.exe WolfProc.R "./input" "../LATEX/Tables"  NoFile R D
rscript.exe WolfProc.R "./input" "../LATEX/Tables"  NoFile R V

if not "%1" == "NoPause" PAUSE
```
