## Supplementary material for "Human-caused wolf mortality persists for years after discontinuation of hunting": WolfProc.R

```
 
#if (!require("dplyr")) {
#  install.packages("dplyr", dependencies = TRUE)
#  library(dplyr)
#}

LINE_SOLID = 1
LINE_DASHED = 2
LINE_DOTTED = 3

#if (!require("tidyverse")) {
#  install.packages("tidyverse", dependencies = TRUE)
#  library(tidyverse)
#}

#if (!require("broom")) {
#  install.packages("broom", dependencies = TRUE)
#  library(broom)
#}

NormalizeDailyHazardVariance <- function(df) 
{
  nLength = length(df$DailyHazardVariance)
  for( i in 1:nLength) { #sometimes hazard is 0, and so is variance. However, there are a lot of radio-days. 
    if( df$DailyHazardVariance[i]==0 ) { #thus, rather than tossing data, having a reasonable estimate of weight of this 0 would be preferable
      # no death but we have the radio-days, try 4 nearest neighbors, and estimate average variance value
      nAveVar = 0
      nStart = i-2;
      if(nStart <1 ) {
        nStart = 1
      }
      nEnd = nStart +2
      if(nEnd > nLength) {
        nEnd = nLength
        nStart = nEnd - 4
      }
      
      nValuesUsed = 0
      for( n in nStart:nEnd){
        if( n != i ) {
          if(df$DailyHazardVariance[n] != 0) {
            nAveVar = nAveVar + df$DailyHazardVariance[n] * df$RadioDays[n] / df$RadioDays[i]
            nValuesUsed = nValuesUsed + 1
          }
        }
      }
      
      df$DailyHazardVariance[i] = nAveVar / nValuesUsed
    }
  }
  varianceNormalized <- df$DailyHazardVariance
  #print(df$DailyHazardVariance)
  varianceNormalized
}

    
SaveRegressionCoefficients <- function(YEAR_RANGE, SUBS, szSrcPath, szDstPath, bRegrWeightVariance) {
  
  nYearRangeLength <- length(YEAR_RANGE)
  szInfoText <- "%\r\n"
  
  szWeightTitle = ""
  szWeightTitleText = ""
  if( bRegrWeightVariance ) {
    szWeightTitle = "weightInvVariance"  
    szWeightTitleText = "weighted by inverse variance"  
  } else {
    szWeightTitle = "weightRadioDays"  
    szWeightTitleText = "weighted by radio-days squared"  
  }

  szFunctionTitle <-paste(YEAR_RANGE[1], "-", YEAR_RANGE[nYearRangeLength], "-", szWeightTitle, "-Table2", sep="")
  szDstFile <- paste(szDstPath,"/", szFunctionTitle, ".txt", sep="")
  
  # save ourselves a trouble of writing a lot of lines for a table preamble, by copying the header file; however the title is missing
  szSrcHeader <- paste("./TableHeader2.txt", sep="")

  #make the function name that takes any letters or numbers, with two parameters
  szText <- paste("\\expandafter\\newcommand\\csname ", szFunctionTitle, "\\endcsname[3] {", sep="")
  cat(szText, file=szDstFile, append=FALSE, sep = "\r\n") #overwrite the file
  
  #now copy the header file, do not overwrite
  szText <- paste(readLines(szSrcHeader), collapse="\n") #read the file, will lose the lasr \r\n
  cat(szText, file=szDstFile, append=TRUE, sep = "\r\n") # append it into destination, adding the now missing new line
  
  szYearStartMonth <- c( "Jan", "Feb", "Mar", "Apr","May","Jun","Jul","Aug","Sep","Oct","Nov","Dec");

  nEntry <- 1
  Entries <- length(szYearStartMonth)
  
  # load summary prevalences for all diseases for further analysis
  while ( nEntry <= Entries) {
    szSrcFile <- paste(szSrcPath, "/", szYearStartMonth[nEntry], ".csv", sep="")
    
    if( file.exists( szSrcFile) ) { 
      df <- read.csv(szSrcFile, header=TRUE, sep=",", stringsAsFactors = FALSE)
      if( bRegrWeightVariance ) {
        df$DailyHazardVariance <- NormalizeDailyHazardVariance(df) 
      }
      if( df$Year[1] == 2004 && df$Year[2] == 2005 ) { # must have valid rows, this proves it
        if( bRegrWeightVariance ) {
          wt <- 1.0 / df$DailyHazardVariance # form weights as a standard inverse of Variance of the Radio-days and wolf deaths
        } else {
          wt <- df$RadioDays^2 #1.0 / df$DailyHazardVariance # form weights proportionate to Radio-days in a period
        }
        
        # run weigted lm()
        fit <- lm(DailyHazard~Year, data = df, subset=SUBS, weights=wt) #weighted
        r2 = summary(fit)$r.squared
        r2adj = summary(fit)$adj.r.squared
        stdErr <-summary(fit)$coefficients[2, 2]
        intercept <- summary(fit)$coefficients[1,1]
        slope = summary(fit)$coefficients[2,1]
        f <- summary(fit)$fstatistic
        p <- pf(f[1],f[2],f[3],lower.tail=F)
        attributes(p) <- NULL
        
        conf_int <- predict(fit, interval="confidence", level = 0.95)
        
        #recalculate mean survival as a product of yearly survivals power 1/years counted
        dbSurvivalMean <- 1.0
        for(nSurv in 1:nYearRangeLength) {
          dbSurvivalMean = dbSurvivalMean * (1.0 - conf_int[nSurv,1])^365
          nSurv = nSurv + 1
        }
        dbSurvivalMean = dbSurvivalMean^(1.0/nYearRangeLength)

        szText <- paste( paste("\t\t\t\t", szYearStartMonth[nEntry]), 
                         format(stdErr, digits=2, nsmall=6, scientific=FALSE), 
                         format(r2, digits=1, nsmall=4, scientific=FALSE), 
                         format(slope, digits=2, nsmall=5, scientific=FALSE), 
                         paste("\\textcolor{txtGray}{", format(intercept, digits=1, nsmall=1), "}"), 
                         format(dbSurvivalMean, digits=3, nsmall=3),
                         format((1.0 - conf_int[1,1])^365, digits=3, nsmall=3), 
                         paste(format((1.0 - conf_int[nYearRangeLength,1])^365, digits=3, nsmall=3), " \\\\"), sep=" & ")
        
        cat(szText, file=szDstFile, append=TRUE, sep = "\r\n")
      }
    }
    nEntry = nEntry + 1
  }  
  cat("\t\t\t\t\\bottomrule", file=szDstFile, append=TRUE, sep = "\r\n")
  cat("\t\t\t\\end{tabular*} \r\n", file=szDstFile, append=TRUE, sep = "")
  szInfoText <- paste( "\t\t\t\\footnotetext[]{Statistics for years: ", YEAR_RANGE[1], " - ", YEAR_RANGE[nYearRangeLength], ", linear regression ", szWeightTitleText, 
            ". 'stdError' is standard error, '$r^2$', 'slope' and 'intercept' of the  linear regression fit. Survival 'begin' and 'end' are survivals in the first and last year of the period.",
                       "  #3}", sep="")
  cat(szInfoText, file=szDstFile, append=TRUE, sep = "\r\n")
  cat("\t\t\\end{minipage} \r\n\t\\end{center} \r\n\\end{table}\r\n} \r\n", file=szDstFile, append=TRUE, sep = "")
}


FillConfidenceIntervals <- function(fit, lwr, upr)
{ # this is a convenient to pass and save the regression parameters
  conf$fit <- fit
  conf$lwr <- lwr
  conf$upr <- upr
  conf
}

  
PerformDailyHazardRegression <- function(df, x, SUBS, bRegrWeightVariance) { # returns 
  # calculate weights
  if( bRegrWeightVariance ) {
    wt <- 1.0 / df$DailyHazardVariance # form weights as a standard inverse of Variance of the Radio-days and wolf deaths
  } else {
    wt <- df$RadioDays^2  # form weights proportionate to Radio-days in a period
  }
  
  # run weigted lm()
  fit <- lm(df[,"DailyHazard"]~Year, data = df, subset=SUBS, weights=wt)
  #  fit <- lm(df[,"DailyHazard"]~Year, data = df, subset=SUBS) #comparison without weights - OF COURSE NEED WEIGHTS!
  conf_interval <- predict(fit, newdata=data.frame(Year=x), interval="confidence", level = 0.95)
  conf_interval # return this ready to use
}

# main drawing of trendlines and confidence intervals
AndDrawLines <- function(conf_interval, x, colLine, colFill, ltY) {
  matlines(x, conf_interval[,2:3], col=colLine, lty=1, lwd = 0.5)
  matlines(x, conf_interval[,1], col=colLine,lty=ltY, lwd = 3)
  polygon(c(x, rev(x)), c(conf_interval[,3], rev(conf_interval[,2])), col=colFill, border=NA)
}

# perform a simple regression of our Dats Frames
PerformRegression <- function(df, ColToRegress, x, SUBS) {
  fit <- lm(df[,ColToRegress]~Year, data = df, subset=SUBS) #comparison without weights
  conf_interval <- predict(fit, newdata=data.frame(Year=x), interval="confidence", level = 0.95)
  conf_interval
}

# prepare a subset of an array
PrepareNewColumnForDataFrame <- function(nYearsInArray, SubsetToInsertInto, DataToPassOver) {
  NewColumn <- rep(NA, nYearsInArray)
  NewColumn[SubsetToInsertInto] <- DataToPassOver
  NewColumn
}

# prepare concatenation of a subset of two arrays
PrepareTwoNewColumnsForDataFrame <- function(nYearsInArray, Subset1toInsertInto, Data1toPassOver, Subset2toInsertInto, Data2toPassOver) {
  NewColumn <- rep(NA, nYearsInArray)
  NewColumn[Subset1toInsertInto] <- Data1toPassOver
  NewColumn[Subset2toInsertInto] <- Data2toPassOver
  NewColumn
}

# First function to start any plot(), and make axex and scatter
PlotHeaderWithScatter <- function(x, y, colorGroup, yLabel, Title, yMin, yMax, yAxisScale, bShowBeforeAfterText = FALSE) {
  plot(x, y, col = colorGroup, lwd = 1.5, xlab="Year", ylab=NA, main=NA, xaxt="n", yaxt="n", ylim = c(yMin,yMax), xlim = c(2004,2019),
       bty = "L",#axes = FALSE,
       cex.main = 0.1,   # Title size
       cex.sub = 1.4,  # Subtitle size
       cex.lab = 1.5,    # X-axis and Y-axis labels size
       cex.axis = 1,
       ) # Axis labels size)
  axis(side=1, at=seq(2004, 2019, by=1), cex.axis=1.4)
  myLabelPos = seq(yMin, yMax, by=yAxisScale)
  if( yAxisScale < 0.01) {
    myLabelText = sprintf(myLabelPos,fmt = '%#.4f')
    axis(side=2, at=myLabelPos, labels=myLabelText, las=1, cex.axis=1.4)    # las=1 Horizontal labels) par(mgp=c(axis.title.position, axis.label.position, axis.line.position))
    title(ylab = yLabel, cex.lab = 1.5,  line = 4.9)
  } else {
    axis(side=2, at=myLabelPos, las=1, cex.axis=1.4)    # las=1 Horizontal labels)
    title(ylab = yLabel, cex.lab = 1.5,  line = 3.0)
  }
  if(bShowBeforeAfterText) {
    text(x = 2007, y = yMax,  "Before hunt (2004-2011)",col=rgb(0, 0.3,0))
    text(x = 2015, y = yMax,  "Following first hunt (2012-2019)",col=rgb(0.6,0, 0))
  }
}


SaveRegressionDiscontinuity <- function(YEAR_RANGE, szSrcPath, szDstPath, bRegrWeightVariance) {
  
  if (!is.element("dplyr", installed.packages()[,1])) {
    install.packages("dplyr", dep = TRUE)
  }
  #require("dplyr", character.only = TRUE)
  library(dplyr)
  
  nYearRangeLength <- length(YEAR_RANGE)
  szInfoText <- "%\r\n"
  
  szWeightTitle = ""
  szWeightTitleText = ""
  if( bRegrWeightVariance ) {
    szWeightTitle = "weightInvVariance"  
    szWeightTitleText = "weighted by inverse variance"  
  } else {
    szWeightTitle = "weightRadioDays"  
    szWeightTitleText = "weighted by radio-days squared"  
  }
  
  szFunctionTitle <- "RegressionDiscontinuity-Table4"
  szDstFile <- paste(szDstPath,"/", szFunctionTitle, ".txt", sep="")
  
  # save ourselves a trouble of writing a lot of lines for a table preamble, by copying the header file; however the title is missing
  szSrcHeader <- paste("./TableHeader4.txt", sep="")
  
  #make the function name that takes any letters or numbers, with two parameters
  szText <- paste("\\expandafter\\newcommand\\csname ", szFunctionTitle, "\\endcsname[3] {", sep="")
  cat(szText, file=szDstFile, append=FALSE, sep = "\r\n") #overwrite the file
  
  #now copy the header file, do not overwrite
  szText <- paste(readLines(szSrcHeader), collapse="\n") #read the file, will lose the lasr \r\n
  cat(szText, file=szDstFile, append=TRUE, sep = "\r\n") # append it into destination, adding the now missing new line
  
  szYearStartMonth <- c( "Jan", "Feb", "Mar", "Apr","May","Jun","Jul","Aug","Sep","Oct","Nov","Dec");
  
  nEntry <- 1
  Entries <- length(szYearStartMonth)
  
  # load summary prevalences for all diseases for further analysis
  while ( nEntry <= Entries) {
    szSrcFile <- paste(szSrcPath, "/", szYearStartMonth[nEntry], ".csv", sep="")
    
    if( file.exists( szSrcFile) ) { 
      df <- read.csv(szSrcFile, header=TRUE, sep=",", stringsAsFactors = FALSE)
      if( bRegrWeightVariance ) {
        df$DailyHazardVariance <- NormalizeDailyHazardVariance(df) 
      }
      if( df$Year[1] == 2004 && df$Year[2] == 2005 ) { # must have valid rows, this proves it
        wt <- 0
        if( bRegrWeightVariance ) {
          wt <- 1.0 / df$DailyHazardVariance # form weights as a standard inverse of Variance of the Radio-days and wolf deaths
        } else {
          wt <- df$RadioDays^2 #1.0 / df$DailyHazardVariance # form weights proportionate to Radio-days in a period
        }
        
        nFiveYearRange = 5
        pValue = rep( NA, nFiveYearRange) 
        nDstep = rep( NA, nFiveYearRange) 
        szDiscText = rep( "", nFiveYearRange) 
        for( nY in 1:nFiveYearRange ) {
          df1 = mutate(df, D = ifelse(df$Year >= (2009 + nY), 1, 0)) #don't need all, years 2010--2014
          fit <- lm(MortalityRaw ~ D * I(Year - (2009 + nY)), data = df1, weights=wt)
          summary(fit)
          pValue[nY] = summary(fit)$coefficients[2,4]
          nDstep[nY] = summary(fit)$coefficients[2,1]
          szStarText = " ("
          if( pValue[nY] <= 0.05 ) {
            szStarText = "\\textbf{*} ("
          }
          szDiscText[nY] = paste(format(pValue[nY], digits=2, nsmall=3, scientific=FALSE), szStarText, 
                                 round(nDstep[nY]*100,0), "\\%)", sep="")
          
          if( nEntry == 11 ) {
            if( nY >= 2 && nY <=4 ) {
              szPdfName <- paste(szDstPath,"/", "RegrDiscont-", szYearStartMonth[nEntry], "-", (2009 + nY), ".PDF", sep="")
              pdf(szPdfName)
              
              par(mar = c(4, 4.4, 0.2, 0.3)) # bottom, left, top, right in line heights
              conf_int <- predict(fit, interval="confidence", level = 0.95)
              PlotHeaderWithScatter(df$Year, df$MortalityRaw, rgb(0,0,0.2), "Yearly Mortality", szFileName, 0.0, 0.7, 0.1, TRUE)
              AndDrawLines( conf_int, YEAR_RANGE, rgb(0.8, 0.6, 0), rgb(0.7, 0.6, 0, 0.15), LINE_SOLID)
              
              dev.off()
            }
          }
        }        
       
        szText <- paste( paste("\t\t\t\t", szYearStartMonth[nEntry], sep = ""), 
                          szDiscText[1], szDiscText[2], szDiscText[3], szDiscText[4], 
                          paste(szDiscText[5], " \\\\", sep = ""), sep=" & ")
        cat(szText, file=szDstFile, append=TRUE, sep = "\r\n")
      }
    }
    nEntry = nEntry + 1
  }  
  cat("\t\t\t\t\\bottomrule", file=szDstFile, append=TRUE, sep = "\r\n")
  cat("\t\t\t\\end{tabular*} \r\n", file=szDstFile, append=TRUE, sep = "")
  cat("\t\t\t #3", file=szDstFile, append=TRUE, sep = "\r\n")
  cat("\t\t\\end{minipage} \r\n\t\\end{center} \r\n\\end{table}\r\n} \r\n", file=szDstFile, append=TRUE, sep = "")
}


#here I perform all analysis and save it into the Figure PDFs and CSVs ready for manuscript and supplementary
main <- function() {
  args <- commandArgs(trailingOnly = TRUE)
  
  bRunManual <- 1 # this is in case I run manually
  
  if( length(args) >= 3 ) { #if run from .BAT file, manual is overwitten to <-0
    bRunManual <- 0
  }

  bSaveRegrStats <- 0
  bRegrWeightVariance <- 0 # do weighing by rDays^2 DEFAULT
  bSaveRegrDiscontinuity <- 0
  
  if( bRunManual == 1 ) { #debugging set of parameters
    setwd("/code/WolfDNR/R")
    szSrcPath <- "./input" #args[1]
    szDstPath <-  "./result" #args[2]  # path to the result file
    szFileName <-  "Sep" #args[3]
    bSaveRegrStats = 1
    bRegrWeightVariance <- 0
  } else { # batch file set of parameters
    szSrcPath <- args[1]
    szDstPath <-  args[2]  # path to the result file
    szFileName <- args[3]
    for( i in 1:length(args)) {
      if( args[i] == "V" ) {
        bRegrWeightVariance = 1 # do weighing by 1/Variance, not rDays^2
      } else if( args[i] == "R" ) {
        bSaveRegrStats = 1  # only save a Table of each month linear regression fit coefficients running SaveRegressionCoefficients 3 times - covering 3 periods
      } else if( args[i] == "D" ) {
        bSaveRegrDiscontinuity = 1
      }
    }
  }

  if (file.exists(szDstPath) == FALSE) { # create destination directory if necessary
    dir.create(szDstPath)
  }

  RANGE_8y = 1:8
  RANGE_16y = 1:16
  
  SUBS_1_16 = 1:16
  SUBS_1_8 = 1:8
  SUBS_9_16 = 9:16
  
  YEARS_2004_2019 = seq(2004,2019,by = 1)
  YEARS_2004_2011 = seq(2004,2011,by = 1)
  YEARS_2012_2019 = seq(2012,2019,by = 1)

  if(  bSaveRegrDiscontinuity ) {
    SaveRegressionDiscontinuity(YEARS_2004_2019, szSrcPath, szDstPath, bRegrWeightVariance) 
  }

  if( bSaveRegrStats ) {
    SaveRegressionCoefficients(YEARS_2004_2019, SUBS_1_16, szSrcPath, szDstPath, bRegrWeightVariance)
    
    SaveRegressionCoefficients(YEARS_2004_2011, SUBS_1_8, szSrcPath, szDstPath, bRegrWeightVariance)
    
    SaveRegressionCoefficients(YEARS_2012_2019, SUBS_9_16, szSrcPath, szDstPath, bRegrWeightVariance)
  }
  
  if( bSaveRegrStats || bSaveRegrDiscontinuity ) {
    return(5)
  }

  szDstFile <- paste(szDstPath,"/", szFileName,  ".csv", sep="")
  szSrcFile <- paste(szSrcPath,"/", szFileName, ".csv", sep="")
  
  df <- read.csv(szSrcFile, header=TRUE, sep=",", stringsAsFactors = FALSE)

  nYearsInArray <- length(df$Year)

  # make sure we have estimates for missing Variance values, else small number of data points will be ignored
  if( bRegrWeightVariance ) {
    df$DailyHazardVariance <- NormalizeDailyHazardVariance(df) 
  }

  # Change the margins of the plot (the first is the bottom margin)
  #par(mar = c(4, 4.2, 0.2, 0.3)) # bottom, left, top, right in line heights
  
  if( bRunManual == 0) { # FIG 1  Daily Unit Hazard ACROSS ALL YEARS
    szPdfName <- paste(szDstPath,"/", szFileName, "-1", ".PDF", sep="")
    pdf(szPdfName)
  }
  #windows(width = 8, height = 8)
  par(mar = c(4, 6.2, 0.2, 0.3)) # bottom, left, top, right in line heights
  PlotHeaderWithScatter(df$Year, df$DailyHazard, rgb(0,0,0.2), "Daily Unit Hazard", szFileName, 0.0, 0.0035, 0.0005)
  print("2")
  #legend(x = "bottomleft", inset = c(-0.12, -0.32), 
  #       legend = c("before hunt", "including and following hunt", "ouline single range regression", "shaded: 95% confidence interval"),
  #       lty = c(LINE_SOLID, LINE_SOLID, LINE_DOTTED, LINE_SOLID ),   col=c(rgb(0, 0.6, 0),rgb(1, 0, 0),rgb(0.8, 0.8, 0.8),"white"),
  #       lwd = 3, box.col = 0, box.lwd = 0.3, xpd = TRUE, cex=1.15)
  ci_HazardAllYears <- PerformDailyHazardRegression(df, YEARS_2004_2019, SUBS_1_16, bRegrWeightVariance)
  AndDrawLines( ci_HazardAllYears, YEARS_2004_2019, rgb(0.5, 0.5, 1.0), rgb(0.5, 0.5, 1.0, 0.15), LINE_SOLID)
    
  ci_survivalAllYears <- (1.0 - ci_HazardAllYears)^365 #get the Survival for free
 
  # save into DataFrame for saving to CSV
  df$GlobSurv_mean <- ci_survivalAllYears[,1]
  df$GlobSurv_2_5 <- ci_survivalAllYears[,3]
  df$GlobSurv_97_5 <- ci_survivalAllYears[,2]

  if( bRunManual == 0 ) {  # FIG 2  Survival ACROSS ALL YEARS
    dev.off()
    szPdfName <- paste(szDstPath,"/", szFileName, "-2", ".PDF", sep="")
    pdf(szPdfName)
  }
  par(mar = c(4, 4.2, 0.2, 0.3)) # bottom, left, top, right in line heights
  PlotHeaderWithScatter(df$Year, df$SurvivalRaw, rgb(0,0,0.2), "Yearly Survival", szFileName, 0.3, 1.0, 0.1)
  #legend(x = "bottomleft", inset = c(-0.12, -0.32), 
  #       legend = c("before hunt", "including and following hunt", "ouline single range regression", "shaded: 95% confidence interval"),
  #       lty = c(LINE_SOLID, LINE_SOLID, LINE_DOTTED, LINE_SOLID ),   col=c(rgb(0, 0.6, 0),rgb(1, 0, 0),rgb(0.8, 0.8, 0.8),"white"),
  #       lwd = 3, box.col = 0, box.lwd = 0.3, xpd = TRUE, cex=1.15)

  AndDrawLines( ci_survivalAllYears, YEARS_2004_2019, rgb(0.5, 0.5, 1.0), rgb(0.5, 0.5, 1.0, 0.15), LINE_SOLID)
  
  if( bRunManual == 0 ) {  # FIG 3  Daily Unit Hazard BEFORE AFTER HUNT
    dev.off()
    szPdfName <- paste(szDstPath,"/", szFileName, "-3", ".PDF", sep="")
    pdf(szPdfName)
  }
  par(mar = c(4, 6.2, 0.2, 0.3)) # bottom, left, top, right in line heights
  PlotHeaderWithScatter(df$Year, df$DailyHazard, rgb(0,0,0.2), "Daily Unit Hazard", szFileName, 0.0, 0.0035, 0.0005, TRUE)
  
  AndDrawLines( ci_HazardAllYears, YEARS_2004_2019, rgb(0.8, 0.8, 0.8), rgb(0, 0, 0, 0.05), LINE_DASHED)

  conf_int1 <- PerformDailyHazardRegression(df, YEARS_2004_2011, SUBS_1_8, bRegrWeightVariance)
  AndDrawLines( conf_int1, YEARS_2004_2011, rgb(0, 0.6, 0), rgb(0, 1, 0, 0.15), LINE_SOLID)
  conf_int2 <- PerformDailyHazardRegression(df, YEARS_2012_2019, SUBS_9_16, bRegrWeightVariance)
  AndDrawLines( conf_int2, YEARS_2012_2019, rgb(1, 0, 0), rgb(1, 0, 0, 0.15), LINE_SOLID)
  
  ci_Surv1_8  <- (1.0 - conf_int1)^365
  ci_Surv9_15  <- (1.0 - conf_int2)^365
  
  df$Surv_mean <- PrepareTwoNewColumnsForDataFrame(nYearsInArray, SUBS_1_8, ci_Surv1_8[,1], SUBS_9_16, ci_Surv9_15[,1])
  df$Surv_2_5  <- PrepareTwoNewColumnsForDataFrame(nYearsInArray, SUBS_1_8, ci_Surv1_8[,3], SUBS_9_16, ci_Surv9_15[,3])
  df$Surv_97_5  <- PrepareTwoNewColumnsForDataFrame(nYearsInArray, SUBS_1_8, ci_Surv1_8[,2], SUBS_9_16, ci_Surv9_15[,2])
  
  if( bRunManual == 0 ) {  # FIG 4  Survival BEFORE AFTER HUNT
    dev.off()
    szPdfName <- paste(szDstPath,"/", szFileName, "-4", ".PDF", sep="")
    pdf(szPdfName)
  }
  par(mar = c(4, 4.2, 0.2, 0.3)) # bottom, left, top, right in line heights
  PlotHeaderWithScatter(df$Year, df$SurvivalRaw, rgb(0,0,0.2), "Yearly Survival", szFileName, 0.3, 1.0, 0.1, TRUE)
  # repeat the all years outline  
  AndDrawLines( ci_survivalAllYears, YEARS_2004_2019, rgb(0.8, 0.8, 0.8), rgb(0, 0, 0, 0.05), LINE_DASHED)
  # draw hazard before and after
  AndDrawLines( ci_Surv1_8, YEARS_2004_2011, rgb(0, 0.6, 0), rgb(0, 1, 0, 0.15), LINE_SOLID)
  AndDrawLines( ci_Surv9_15, YEARS_2012_2019, rgb(1, 0, 0), rgb(1, 0, 0, 0.15), LINE_SOLID)
  
  if( bRunManual == 0 ) {  # FIG 5 now show mortality BEFORE AFTER HUNT
    dev.off()
    szPdfName <- paste(szDstPath,"/", szFileName, "-5", ".PDF", sep="")
    pdf(szPdfName)
  }
  mortality_ci1  <- (1.0 - ci_Surv1_8)
  mortality_ci2  <- (1.0 - ci_Surv9_15)
  
  par(mar = c(4, 4.2, 0.2, 0.3)) # bottom, left, top, right in line heights
  PlotHeaderWithScatter(df$Year, df$MortalityRaw, rgb(0,0,0.2), "Yearly Mortality", szFileName, 0.0, 0.7, 0.1, TRUE)
  AndDrawLines( mortality_ci1, YEARS_2004_2011, rgb(0.8, 0.6, 0), rgb(0.7, 0.6, 0, 0.15), LINE_SOLID)
  AndDrawLines( mortality_ci2, YEARS_2012_2019, rgb(0.8, 0.6, 0), rgb(0.7, 0.6, 0, 0.15), LINE_SOLID)
  

  if( bRunManual == 0 ) {  # FIG 6 calculate natural and human causes BEFORE AFTER HUNT and draw
    dev.off()
    szPdfName <- paste(szDstPath,"/", szFileName, "-6", ".PDF", sep="")
    pdf(szPdfName)
  }

  # calculate  the fraction of mortality by cause based on daily hazard ratio for each year  
  MortalityHuman <- rep( NA, nYearsInArray)
  MortalityNatural <- rep( NA, nYearsInArray)
  for( nPos in 1:nYearsInArray) {
    if( df$DailyHazard[nPos] > 0.0000001 ) {
      MortalityHuman[nPos] = df$DailyHazardHuman[nPos] * (1.0 - df$Surv_mean[nPos]) / df$DailyHazard[nPos]
      MortalityNatural[nPos] = df$DailyHazardNatural[nPos] * (1.0 - df$Surv_mean[nPos]) / df$DailyHazard[nPos]
    }
  }

  # save in DF to save in CSV
  df$MortalityHuman <- MortalityHuman
  df$MortalityNatural <- MortalityNatural

  # produce double set - Natural and Human - for scatter plot
  newData <- c(MortalityHuman, MortalityNatural) # data itself
  newYear <- c(YEARS_2004_2019, YEARS_2004_2019) # with corresponding years
  newGroup <- c(rep(rgb(0.8,0,0), 16) , rep(rgb(0,0.5,0), 16)) # and colors of the data points
                
  par(mar = c(4, 4.2, 0.2, 0.3)) # bottom, left, top, right in line heights
  PlotHeaderWithScatter(newYear, newData, newGroup, "Yearly Mortality", szFileName, 0.0, 0.7, 0.1, TRUE) # the scatter plot itself

  AndDrawLines( mortality_ci1, YEARS_2004_2011, rgb(0.0, 0.0, 0), rgb(0.7, 0.6, 0, 0.0), LINE_DASHED)
  AndDrawLines( mortality_ci2, YEARS_2012_2019, rgb(0.0, 0.0, 0), rgb(0.8, 0.6, 0, 0.0), LINE_DASHED)

  ci_nat_1 <- PerformRegression(df, "MortalityNatural", YEARS_2004_2011, SUBS_1_8) 
  ci_nat_2 <- PerformRegression(df, "MortalityNatural", YEARS_2012_2019, SUBS_9_16) 
  AndDrawLines( ci_nat_1, YEARS_2004_2011, rgb(0, 0.6, 0), rgb(0, 1, 0, 0.15), LINE_SOLID)
  AndDrawLines( ci_nat_2, YEARS_2012_2019, rgb(0, 0.6, 0), rgb(0, 1, 0, 0.15), LINE_SOLID)
  
  
  ci_hum_1 <- PerformRegression(df, "MortalityHuman", YEARS_2004_2011, SUBS_1_8) 
  ci_hum_2 <- PerformRegression(df, "MortalityHuman", YEARS_2012_2019, SUBS_9_16) 
  AndDrawLines( ci_hum_1, YEARS_2004_2011, rgb(1, 0, 0), rgb(1, 0, 0, 0.15), LINE_SOLID)
  AndDrawLines( ci_hum_2, YEARS_2012_2019, rgb(1, 0, 0), rgb(1, 0, 0, 0.15), LINE_SOLID)
  
  if( bRunManual == 0 ) { # close complete figures
    dev.off()
  }
  
  
  df$MortNatural_mean <- PrepareTwoNewColumnsForDataFrame(nYearsInArray, SUBS_1_8, ci_nat_1[,1], SUBS_9_16, ci_nat_2[,1])
  df$MortNatural_2_5  <- PrepareTwoNewColumnsForDataFrame(nYearsInArray, SUBS_1_8, ci_nat_1[,2], SUBS_9_16, ci_nat_2[,2])
  df$MortNatural_97_5  <- PrepareTwoNewColumnsForDataFrame(nYearsInArray, SUBS_1_8, ci_nat_1[,3], SUBS_9_16, ci_nat_2[,3])

  df$MortHuman_mean <- PrepareTwoNewColumnsForDataFrame(nYearsInArray, SUBS_1_8, ci_hum_1[,1], SUBS_9_16, ci_hum_2[,1])
  df$MortHuman_2_5  <- PrepareTwoNewColumnsForDataFrame(nYearsInArray, SUBS_1_8, ci_hum_1[,2], SUBS_9_16, ci_hum_2[,2])
  df$MortHuman_97_5  <- PrepareTwoNewColumnsForDataFrame(nYearsInArray, SUBS_1_8, ci_hum_1[,3], SUBS_9_16, ci_hum_2[,3])

  
  { 
    #these two are raw mortalities. They are needed for PerformRegressionAndDrawLines(), but redundant in the file, and thus confusing. Will delete them before saving the file
    df <- subset (df, select = -c(MortalityHuman,MortalityNatural) )
    #save it all to the Result file
    write.table(format( df, digits=6, scientific=FALSE, trim = TRUE),
                file = szDstFile,
                row.names = FALSE,
                col.names = TRUE,
                sep = ",",
                quote = FALSE )  
  }

}

# CALL MY MAIN() 
main()
  
```
