## Supplementary material for "Human-caused wolf mortality persists for years after discontinuation of hunting": WolfSummaryTableBeforeAfter.R

```
PtxtG <- function(value) {
  szText <- paste(format(round(value*100,1), nsmall=1), "\\%", sep="")
  szText
}

DtxtG <- function(value) {
  szText <- paste(format(round(value,2), nsmall=2), sep="")
  szText
}

MakeSurvivalText <- function(df, nRow) {
  szText <- paste(format(round(df$SurvivalRaw[nRow],3), nsmall=3), " (", 
                  format(round(df$ciLwrSmr[nRow],2), nsmall=2), "-", 
                  format(round(df$ciUprSmr[nRow],2), nsmall=2), ")", sep="")
  szText
}

main <- function() {
  args <- commandArgs(trailingOnly = TRUE)

  bRunManual <- 1
  
  if( length(args) >= 3 ) {
    bRunManual <- 0
  }

  if( bRunManual == 1 ) {
    setwd("/code/WolfDNR/R")
    szSrcPath <- "./summaries" #args[1]
    szDstPath <-  "./result" #args[2]  # path to the result file
    szFileName <-  "Nov" #args[3]
  } else {
    szSrcPath <- args[1]
    szDstPath <-  args[2]  # path to the result file
    szFileName <- args[3]
  }

  if (file.exists(szDstPath) == FALSE) { # create destination directory if necessary
    dir.create(szDstPath)
  }
  
  szFunctionTitle <-paste(szFileName,  "-Table1", sep="")
  szDstFile <- paste(szDstPath,"/", szFunctionTitle, ".txt", sep="")

  # save ourselves a trouble of writing a lot of lines in table preamble
  szSrcHeader <- paste("./TableHeader1.txt", sep="")
  
  #make the function name that takes any letters or numbers, with two parameters
  szText <- paste("\\expandafter\\newcommand\\csname ", szFunctionTitle, "\\endcsname[3] {", sep="")
  cat(szText, file=szDstFile, append=FALSE, sep = "\r\n") #overwrite the file
  
  #now copy the header file, do not overwrite
  szText <- paste(readLines(szSrcHeader), collapse="\n") #read the file, will lose the lasr \r\n
  cat(szText, file=szDstFile, append=TRUE, sep = "\r\n") # append it into destination, adding the now missing new line

  fileAppendix <- c( ".csv", "a.csv", "M.csv", "F.csv", "y.csv" );
  sectionTitle <- c( "All wolves", "Adult", "Male", "Female", "Juvenile" );

  nEntry <- 1
  Entries <- length(fileAppendix)

  # load summary prevalences for all diseases for further analysis
  while ( nEntry <= Entries) {
    szSrcFile <- paste(szSrcPath, "/", szFileName, fileAppendix[nEntry], sep="")
    
    df <- read.csv(szSrcFile, header=TRUE, sep=",", stringsAsFactors = FALSE)
    if( df$Year[1] == 2011 && df$Year[2] == 2012 ) { # must have at least two valid rows, this proves it
      szText <- paste("\t\t\t\t\\multicolumn{7}{@{}c@{}}{\\textit{", sectionTitle[nEntry], "}}	\\\\" , sep="")
      cat(szText, file=szDstFile, append=TRUE, sep = "\r\n")
      cat("\t\t\t\t\\midrule", file=szDstFile, append=TRUE, sep = "\r\n")
      
      szText <- paste("\t\t\t\t\\textcolor{txtG}{\\bf Before}", MakeSurvivalText(df, 1), PtxtG(df$MortalityRaw[1]), PtxtG(df$MortalityHumanSmr[1]),  PtxtG(df$MortalityNaturalSmr[1]), df$RadioDays[1],  DtxtG(df$DeadHuman[1]),  paste(DtxtG(df$DeadNatural[1]), "\\\\"), sep=" & ")
      cat(szText, file=szDstFile, append=TRUE, sep = "\r\n")
  
      szText <- paste("\t\t\t\t\\textcolor{txtR}{\\bf After}", MakeSurvivalText(df, 2), PtxtG(df$MortalityRaw[2]), PtxtG(df$MortalityHumanSmr[2]),  PtxtG(df$MortalityNaturalSmr[2]),  df$RadioDays[2],  DtxtG(df$DeadHuman)[2],  paste(DtxtG(df$DeadNatural[2]), "\\\\"), sep=" & ")
      cat(szText, file=szDstFile, append=TRUE, sep = "\r\n")

      szText <- paste("\t\t\t\t All", MakeSurvivalText(df, 3), PtxtG(df$MortalityRaw[3]), PtxtG(df$MortalityHumanSmr[3]), PtxtG(df$MortalityNaturalSmr[3]),  df$RadioDays[3],  DtxtG(df$DeadHuman)[3],  paste(DtxtG(df$DeadNatural[3]), "\\\\"), sep=" & ")
      cat(szText, file=szDstFile, append=TRUE, sep = "\r\n")
      
      cat("\t\t\t\t\\bottomrule", file=szDstFile, append=TRUE, sep = "\r\n")
    }
    nEntry = nEntry + 1
  }  
  #cat("\t\t\t\\end{tabular*} \r\n\t\t\\end{minipage} \r\n\t\\end{center} \r\n\\end{table}\r\n}\r\n", file=szDstFile, append=TRUE, sep = "")
  
  cat("\t\t\t\\end{tabular*} \r\n", file=szDstFile, append=TRUE, sep = "")
  cat("\t\t\t #3", file=szDstFile, append=TRUE, sep = "\r\n")
  cat("\t\t\\end{minipage} \r\n\t\\end{center} \r\n\\end{table}\r\n} \r\n", file=szDstFile, append=TRUE, sep = "")
  
}

# CALL MY MAIN() 
main()
  
```
