## Supplementary material for "Human-caused wolf mortality persists for years after discontinuation of hunting": WolfTableByYear.R

```
PtxtG <- function(value) {
  szText <- paste(format(round(value*100,1), nsmall=1), "\\%", sep="")
  szText
}

DtxtG <- function(value) {
  szText <- paste(format(round(value,2), nsmall=2), sep="")
  szText
}


main <- function() {
  args <- commandArgs(trailingOnly = TRUE)

  bRunManual <- 1
  
  if( length(args) >= 3 ) {
    bRunManual <- 0
  }

  if( bRunManual == 1 ) {
    setwd("/code/WolfDNR/R")
    szSrcPath <- "./summaries" #args[1]
    szDstPath <-  "./result" #args[2]  # path to the result file
    szFileName <-  "Nov" #args[3]
  } else {
    szSrcPath <- args[1]
    szDstPath <-  args[2]  # path to the result file
    szFileName <- args[3]
  }

  if (file.exists(szDstPath) == FALSE) { # create destination directory if necessary
    dir.create(szDstPath)
  }
  
  szFunctionTitle <-paste(szFileName,  "-Table3", sep="")
  szDstFile <- paste(szDstPath,"/", szFunctionTitle, ".txt", sep="")

  # save ourselves a trouble of writing a lot of lines in table preamble
  szSrcHeader <- paste("./TableHeader3.txt", sep="")
  
  #make the function name that takes any letters or numbers, with two parameters
  szText <- paste("\\expandafter\\newcommand\\csname ", szFunctionTitle, "\\endcsname[3] {", sep="")
  cat(szText, file=szDstFile, append=FALSE, sep = "\r\n") #overwrite the file
  
  #now copy the header file, do not overwrite
  szText <- paste(readLines(szSrcHeader), collapse="\n") #read the file, will lose the lasr \r\n
  cat(szText, file=szDstFile, append=TRUE, sep = "\r\n") # append it into destination, adding the now missing new line

  szSrcFile <- paste(szSrcPath, "/", szFileName, ".csv", sep="")
  
  df <- read.csv(szSrcFile, header=TRUE, sep=",", stringsAsFactors = FALSE)
  

  nEntry <- 1
  nYearsInArray <- length(df$Year)

  # load summary prevalences for all diseases for further analysis
  for( nPos in 1:nYearsInArray) {

    szText <- paste( paste("\t\t\t\t ", df$Year[nPos], sep=""), df$RadioDays[nPos], df$Dead[nPos], 
                           format(df$DeadHuman[nPos], digits=3, nsmall = 3), format(df$DeadNatural[nPos], digits=3, nsmall = 3), 
                           paste("\\textcolor{txtGray}{", format(df$DailyHazard[nPos], digits=3, nsmall = 3, scientific=TRUE), "}"), 
                           paste(format(df$DailyHazardVariance[nPos], digits=3, nsmall = 3, scientific=TRUE), 
                  " \\\\"), sep=" & ")
    cat(szText, file=szDstFile, append=TRUE, sep = "\r\n")
  }  
  cat("\t\t\t\t\\bottomrule", file=szDstFile, append=TRUE, sep = "\r\n")
  cat("\t\t\t\\end{tabular*} \r\n", file=szDstFile, append=TRUE, sep = "")
  cat("\t\t\t #3", file=szDstFile, append=TRUE, sep = "\r\n")
  cat("\t\t\\end{minipage} \r\n\t\\end{center} \r\n\\end{table}\r\n} \r\n", file=szDstFile, append=TRUE, sep = "")
  
}

# CALL MY MAIN() 
main()
  
```
