## Supplementary material for "Human-caused wolf mortality persists for years after discontinuation of hunting": RunExe.bat

```
".\WolfDnr.exe"  ".\MnWolfSurvivalMortalityAge.csv" ".\R\input" 12 W
".\WolfDnr.exe"  ".\MnWolfSurvivalMortalityAge.csv" ".\R\input" 11 W
".\WolfDnr.exe"  ".\MnWolfSurvivalMortalityAge.csv" ".\R\input" 10 W
".\WolfDnr.exe"  ".\MnWolfSurvivalMortalityAge.csv" ".\R\input" 9  W
".\WolfDnr.exe"  ".\MnWolfSurvivalMortalityAge.csv" ".\R\input" 8  W
".\WolfDnr.exe"  ".\MnWolfSurvivalMortalityAge.csv" ".\R\input" 7 W
".\WolfDnr.exe"  ".\MnWolfSurvivalMortalityAge.csv" ".\R\input" 6 W
".\WolfDnr.exe"  ".\MnWolfSurvivalMortalityAge.csv" ".\R\input" 5  W
".\WolfDnr.exe"  ".\MnWolfSurvivalMortalityAge.csv" ".\R\input" 4  W
".\WolfDnr.exe"  ".\MnWolfSurvivalMortalityAge.csv" ".\R\input" 3 W
".\WolfDnr.exe"  ".\MnWolfSurvivalMortalityAge.csv" ".\R\input" 2 W
".\WolfDnr.exe"  ".\MnWolfSurvivalMortalityAge.csv" ".\R\input" 1  W

".\WolfDnr.exe"  ".\MnWolfSurvivalMortalityAge.csv" ".\R\input" 12 N W
".\WolfDnr.exe"  ".\MnWolfSurvivalMortalityAge.csv" ".\R\input" 11 N W
".\WolfDnr.exe"  ".\MnWolfSurvivalMortalityAge.csv" ".\R\input" 10 N W
".\WolfDnr.exe"  ".\MnWolfSurvivalMortalityAge.csv" ".\R\input" 9  N W
".\WolfDnr.exe"  ".\MnWolfSurvivalMortalityAge.csv" ".\R\input" 8  N W
".\WolfDnr.exe"  ".\MnWolfSurvivalMortalityAge.csv" ".\R\input" 7 N W
".\WolfDnr.exe"  ".\MnWolfSurvivalMortalityAge.csv" ".\R\input" 6 N W
".\WolfDnr.exe"  ".\MnWolfSurvivalMortalityAge.csv" ".\R\input" 5  N W
".\WolfDnr.exe"  ".\MnWolfSurvivalMortalityAge.csv" ".\R\input" 4  N W
".\WolfDnr.exe"  ".\MnWolfSurvivalMortalityAge.csv" ".\R\input" 3 N W
".\WolfDnr.exe"  ".\MnWolfSurvivalMortalityAge.csv" ".\R\input" 2 N W
".\WolfDnr.exe"  ".\MnWolfSurvivalMortalityAge.csv" ".\R\input" 1  N W

if not "%1" == "NoPause" PAUSE
```
