## Supplementary material for "Human-caused wolf mortality persists for years after discontinuation of hunting": RunSummaries.bat

```
".\WolfDnr.exe"  ".\MnWolfSurvivalMortalityAge.csv" ".\R\summaries" 11 S W
".\WolfDnr.exe"  ".\MnWolfSurvivalMortalityAge.csv" ".\R\summaries" 11 S y W
".\WolfDnr.exe"  ".\MnWolfSurvivalMortalityAge.csv" ".\R\summaries" 11 S a W
".\WolfDnr.exe"  ".\MnWolfSurvivalMortalityAge.csv" ".\R\summaries" 11 S M W
".\WolfDnr.exe"  ".\MnWolfSurvivalMortalityAge.csv" ".\R\summaries" 11 S F W

".\WolfDnr.exe"  ".\MnWolfSurvivalMortalityAge.csv" ".\R\summaries" 11 S N W
".\WolfDnr.exe"  ".\MnWolfSurvivalMortalityAge.csv" ".\R\summaries" 11 S y N W
".\WolfDnr.exe"  ".\MnWolfSurvivalMortalityAge.csv" ".\R\summaries" 11 S a N W
".\WolfDnr.exe"  ".\MnWolfSurvivalMortalityAge.csv" ".\R\summaries" 11 S M N W
".\WolfDnr.exe"  ".\MnWolfSurvivalMortalityAge.csv" ".\R\summaries" 11 S F N W

if not "%1" == "NoPause" PAUSE
```
