## Supplementary material for "Human-caused wolf mortality persists for years after discontinuation of hunting": stdafx.cpp

```
// stdafx.cpp : source file that includes just the standard includes
// AppentTPed.pch will be the pre-compiled header
// stdafx.obj will contain the pre-compiled type information

#include "stdafx.h"

// TODO: reference any additional headers you need in STDAFX.H
// and not in this file
```
