## Supplementary material for "Human-caused wolf mortality persists for years after discontinuation of hunting": stdafx.h

```
// stdafx.h : include file for standard system include files,
// or project specific include files that are used frequently, but
// are changed infrequently
//

#pragma once

#include "targetver.h"
#include <windows.h>
#include <windowsx.h>
#include <stdio.h>
#include <tchar.h>
/*
class CLocalMemoryObject
{
public:
	void * m_pMemory;
	__int64 m_iMemorySize;
	CLocalMemoryObject( DWORD dAllocationType, __int64 iMemorySize )
	{
		m_iMemorySize = iMemorySize;
		if( iMemorySize ) {
			m_pMemory = GlobalAllocPtr( dAllocationType, m_iMemorySize );
		} else {
			m_pMemory = NULL;
		}
	};
	~CLocalMemoryObject() 
	{
		if( m_pMemory ) {
			GlobalFreePtr( m_pMemory );
			m_pMemory = NULL;
		}
	};
};
*/

// TODO: reference additional headers your program requires here
```
