## Supplementary material for "Human-caused wolf mortality persists for years after discontinuation of hunting": targetver.h

```
#pragma once

// Including SDKDDKVer.h defines the highest available Windows platform.

// If you wish to build your application for a previous Windows platform, include WinSDKVer.h and
// set the _WIN32_WINNT macro to the platform you wish to support before including SDKDDKVer.h.

#include <SDKDDKVer.h>
```
