## Supplementary material for "Human-caused wolf mortality persists for years after discontinuation of hunting": WolfDNR.cpp

```
// AppentTPed.cpp : Defines the entry point for the console application.
//

#include "stdafx.h"
#include <iostream>
#include <direct.h>

using namespace std;

#include "Header.h"

typedef struct RSPatternStruct {
	char szRS[29];
	char cReference;
	char cAlleleSeq;
	BYTE bEntryUsed;
	DWORD dwChromosome;
	DWORD dwPosition;
} RSPatternStruct;
#define MAX_SNPs	100000

BOOL ReadFileInMemBuffer( LPSTR szArgvFileName, BYTE **	ppbFileBfr, __int64 * piFileSize );
__inline BOOL GetTextLine( BYTE * pbFileBfr, __int64 * pnBufPos, char * szTempBuf, int nTempBufSize, __int64 iSrcFileSize );
DWORD CountTextLines( BYTE * pbFileBfr, __int64 iFileSize );
void CreateDir( char * szPath );
double CalcYearlySurvConfIntervals( double dbDailySurv, double dbWolfDays, double dbDeadCount, double * pdbLower, double * pdbUpper );

#define N_YEARS			16
#define N_DAYS_IN_YEAR	365
#define N_MAX_ENTRIES ((N_YEARS +1)*N_DAYS_IN_YEAR)  // +1 because we have one wolf streatching into August 2020, and we will crash if not add one more year

typedef struct AliveDeadStruct {
	DWORD dAlive;
	char cReason[4];
} AliveDeadStruct;

typedef struct MortalityStruct {
	double dbDeadWolvesNatural;
	double dbDeadWolvesHuman;
} MortalityStruct;

typedef struct MortalityOccuredStruct {
	DWORD dYear;
	DWORD dReserved;
	double dbDeadWolvesNatural;
	double dbDeadWolvesHuman;

} MortalityOccuredStruct;

const DWORD dDaysInMonth[14] = { 0,  0,     31,    59,    90,   120,  151,  181,  212,  243,  273,  304,  334,  365 };
const LPSTR szYearStart[13] = { "", "Jan", "Feb", "Mar", "Apr","May","Jun","Jul","Aug","Sep","Oct","Nov","Dec" };


LPSTR szMortCause[4] = { "", "mHuman", "mNatural", "mAll"};

#define AGE_JUVENILE	1
#define AGE_ADULT		2
#define AGE_ALL			(AGE_JUVENILE | AGE_ADULT)
LPSTR szMortAge[4] = { "", "-aJuvenile", "-aAdult", "-aAll"};
char cSelectByAGE = 0, cSelectBySEX = 0;
BOOL cOnlySUMMARY = FALSE;
char   cLastAliveCondition=0;//        varScoreVAF     varScoreEAF     varQuality      hapLink xRef

DWORD YMDtoSequentialDay(DWORD y, DWORD m, DWORD d)
{
	if( y == 0 || m == 0 ) {
		return 0;
	}
	return ( (y - 2004) * 365 + dDaysInMonth[m] + d ); // not accounting for leap years, as it adds 
}

DWORD CountAliveWolfDays(AliveDeadStruct * pAliveDead, DWORD dDay )
{
	DWORD dSumDays = 0;
	DWORD dDayMax = dDay + N_DAYS_IN_YEAR;
	for(DWORD d=dDay; d < dDayMax; d++) {
		dSumDays += pAliveDead[d].dAlive;
	}

	return dSumDays;
}

double CalcMaximumLikelihoodSurvivalRate( double dbWolfDays, double dbDeadCount );
int main( int argc, char *argv[] )
{
	char szDstFile[MAX_PATH];
	char szTempBuf[512] = "", szRsBuf[512] = "", szAddAlleleBfr[16], szID[32], szDeathReason[32], szPlace[32], szAge[32];
	DWORD dStart_Day, dStart_Month, dStart_Year;
	DWORD dEnd_Day, dEnd_Month, dEnd_Year;
	char cSex;
	int nFileCount = 0;
	BYTE *	pbSrcFileBfr, *pbPatternFileBfr;
	__int64 iSrcFileSize = 0, iPatternFileSize = 0;

	DWORD dwLocus, dwPloidy, dwAllele, dwChromosome, dwEntryBeginPos, dwEntryEndPos;


	CLocalMemoryObject cMemArray( GPTR, (__int64)((N_MAX_ENTRIES + N_DAYS_IN_YEAR) * sizeof( AliveDeadStruct )) );
	AliveDeadStruct * pAliveDead = (AliveDeadStruct *)cMemArray.m_pMemory;

	CLocalMemoryObject cMemArray1( GPTR, (__int64)((N_MAX_ENTRIES + N_DAYS_IN_YEAR) * sizeof( MortalityStruct )) );
	MortalityStruct * pMortality = (MortalityStruct *)cMemArray1.m_pMemory;

	CLocalMemoryObject cMemArray2( GPTR, (__int64)((N_MAX_ENTRIES + N_DAYS_IN_YEAR) * sizeof( MortalityOccuredStruct )) );
	MortalityOccuredStruct * pMortalityOccured = (MortalityOccuredStruct *)cMemArray2.m_pMemory;


	if( argc < 4 ) {
		cout << "Incorrect command line. Use: WolfDNR.exe [Path\\SrcFile] [Path\\DstFile] nCause nAge nMonthStart\n";
		return 0;
	}


	if( !ReadFileInMemBuffer( (LPSTR)argv[1], &pbSrcFileBfr, &iSrcFileSize ) ) {  // Read Variation File
		return 0;
	}

	int nYearStartMonth = atoi( argv[3] );
	LPSTR szUnnownsImputation[2] = { "N", "" }; // No Unknowns
	BOOL bImputeUnknowns = 1;
	// here we allow for variable number of letter inputs
	for( int i = 4; i < argc; i++ ) {
		if( argv[i][0] == 'N' ) {
			bImputeUnknowns = 0;
		} else if( argv[i][0] == 'a' || argv[i][0] == 'y' ) {
			cSelectByAGE = argv[i][0];
		} else if( argv[i][0] == 'M' || argv[i][0] == 'F' ) {
			cSelectBySEX = argv[i][0];
		} else if( argv[i][0] == 'S' ) {
			cOnlySUMMARY = argv[i][0];
		} else if( argv[i][0] == 'L' ) {
			cLastAliveCondition = argv[i][0];
		}
	}

	if( cSelectBySEX ) {
		cSelectByAGE = 'a'; // if selecting by sex, do it only for adults
	}

	//_snprintf( szDstFile, sizeof( szDstFile ), "%s%s", argv[2], szUnnownsImputation[bImputeUnknowns] );
	CreateDir( argv[2] );

	DWORD dwBytesWritten;
	DWORD dBeforeHuman = 0, dBeforeNatural = 0, dAfteHuman = 0, dAfterNatural = 0;
	DWORD dHuntCountStartsDate = YMDtoSequentialDay( 2012, nYearStartMonth, 1 ) - dDaysInMonth[nYearStartMonth];

	DWORD dwStartPos = 0;
	DWORD dwPrevVariantPosition = 0;
	DWORD dwCumulativeDays = 0;
	int nFieldsRead = 0;

	// parse the Variation file line by line
	// this is the mai loop where we aggregatein a single  all wolf days, deaths and their causes, and select by age and sex
	for( __int64 iPos = 0; iPos < iSrcFileSize && iPos < iSrcFileSize; iPos++ ) {
		if( !GetTextLine( pbSrcFileBfr, &iPos, szTempBuf, sizeof( szTempBuf ), iSrcFileSize ) ) {
			break;
		}

		DWORD dLastAlive_Month = 0, dLastAlive_Day = 0, dLastAlive_Year = 0;

		if( (nFieldsRead = sscanf( szTempBuf, "%[^,],%[^,],%u/%u/%u,%u/%u/%u,%[^,],%[^,],%c,%u/%u/%u",
			szID, szDeathReason,
			&dStart_Month, &dStart_Day, &dStart_Year, &dEnd_Month, &dEnd_Day, &dEnd_Year,
			szPlace, szAge, &cSex, &dLastAlive_Month, &dLastAlive_Day, &dLastAlive_Year )) >= 10 )
		{
			DWORD dStart = YMDtoSequentialDay( dStart_Year, dStart_Month, dStart_Day );
			DWORD dEnd = YMDtoSequentialDay( dEnd_Year, dEnd_Month, dEnd_Day );
			DWORD dLastAlive = YMDtoSequentialDay( dLastAlive_Year, dLastAlive_Month, dLastAlive_Day );
			DWORD dDaysToAdult = 0;

			if( cLastAliveCondition && dLastAlive > 0 ) {
				if( cLastAliveCondition == 'L' ) {
					dEnd = dLastAlive;
				}
			}

			if( szAge[0] == '2' ) { // there are some entries that say '2' or '2.5' instead of 'adult' they are by definition adults
				szAge[0] = 'a';
			}

			if( cSelectBySEX ) { // 
				if( cSex != cSelectBySEX ) {
					continue;
				}
			}

			if( cSelectByAGE ) { //  there are only about 10000 young days spread over 15 years, not very confident, and yet removing this  interval from adults makes it much more noisy too. Should stick to all wolves aanlysis
				if( szAge[0] != 'a' ) { // if need to account for adult or pup 
					if( dStart_Month < 4 || (dStart_Month == 4 && dStart_Day < 15) ) {
						dDaysToAdult = YMDtoSequentialDay( dStart_Year, 4, 15 ) - dStart;
					} else {
						dDaysToAdult = 365 - (dStart - YMDtoSequentialDay( dStart_Year, 4, 15 ));
					}
					if( szAge[0] == 'p' ) { // if juvenile - done, if pup needs another year
						if( dDaysToAdult < 306 ) { // there are a few pups that seem to be collared at a few days of age, zoologists must have misqualified a yearling as a pup
							dDaysToAdult += 365;
						}
					}
				}
				if( cSelectByAGE == 'a' ) {
					if( szAge[0] != 'a' ) {
						dStart += dDaysToAdult+1;
					}
				} else { // else, limiting age of a pup or yearling to until they mature
					if( dDaysToAdult > 0 ) {
						if( dStart + dDaysToAdult < dEnd ) {
							szDeathReason[0] = 'N'; // obviuosly, the pup survived past this day, it got sequestered
							dEnd = dStart + dDaysToAdult;
						}
					}
				}
				if( (int)dEnd < (int)dStart ) {
					continue;
				}
				if( cSelectByAGE == 'a' && szAge[0] != 'a' ) { // needed adult and matured into adult, can't go other way
					szAge[0] = 'a';
				}
			}

			if( cSelectBySEX ) { // when selecting for sex, discount the pups and juveniles
				if( cSex != cSelectBySEX || szAge[0] != 'a' ) {
					continue;
				}
			}

			dStart -= dDaysInMonth[nYearStartMonth];
			dEnd -= dDaysInMonth[nYearStartMonth];

			if( (int)dStart < 0 ) {
				dStart = 0;  // this is because the offset may be pushing it below Jan 1 2004
			}

			if( !cSelectByAGE || ((cSelectByAGE == 'a' && szAge[0] == 'a') || (cSelectByAGE == 'y' && szAge[0] != 'a')) ) {
				for( DWORD dwPos = dStart; dwPos <= dEnd; dwPos++ ) {
					pAliveDead[dwPos].dAlive += 1;
					dwCumulativeDays++;
				}

				if( szDeathReason[0] != 'N' ) {
					if( pAliveDead[dEnd].cReason[0] == 0 ) {
						pAliveDead[dEnd].cReason[0] = szDeathReason[0];
					} else if( pAliveDead[dEnd].cReason[1] == 0 ) {
						pAliveDead[dEnd].cReason[1] = szDeathReason[0]; // can do it up to 4 entries
					} else if( pAliveDead[dEnd].cReason[2] == 0 ) {
						pAliveDead[dEnd].cReason[2] = szDeathReason[0]; // can do it up to 4 entries
					} else if( pAliveDead[dEnd].cReason[3] == 0 ) {
						pAliveDead[dEnd].cReason[3] = szDeathReason[0]; // can do it up to 4 entries
					}
					if( dEnd >= dHuntCountStartsDate ) {  // find human and natural death count before and after first hunt
						if( szDeathReason[0] == 'h' ) {
							dAfteHuman++;
						} else if( szDeathReason[0] == 'n' ) {
							dAfterNatural++;
						}
					} else {
						if( szDeathReason[0] == 'h' ) {
							dBeforeHuman++;
						} else if( szDeathReason[0] == 'n' ) {
							dBeforeNatural++;
						}
					}
				}
			}
		} else {
			nFieldsRead = nFieldsRead; // just for debugging purposes; we read the entries by position, and the title line will be automatically dropped here
		}
	}
	dwCumulativeDays = dwCumulativeDays;
	const double IMPUTE_BEFORE_HUNT_HUMAN = (double)dBeforeHuman / (double)(dBeforeNatural + dBeforeHuman);
	const double IMPUTE_BEFORE_HUNT_NATURAL = (double)dBeforeNatural / (double)(dBeforeNatural + dBeforeHuman);

	const double IMPUTE_AFTER_HUNT_HUMAN = (double)dAfteHuman / (double)(dAfterNatural + dAfteHuman);
	const double IMPUTE_AFTER_HUNT_NATURAL = (double)dAfterNatural / (double)(dAfterNatural + dAfteHuman);

	// accumulate the cause of death, and impute the 'u' - unknowns - unless specified not to (default is to impute)
	for( DWORD dwPos = 0; dwPos < N_MAX_ENTRIES; dwPos++ ) {
		if( pAliveDead[dwPos].cReason[0] != 0 ) {
			for( DWORD d = 0; d < 4; d++ ) {
				if( pAliveDead[dwPos].cReason[d] == 'h' ) {
					pMortality[dwPos].dbDeadWolvesHuman += 1.0;
				} else if( pAliveDead[dwPos].cReason[d] == 'n' ) {
					pMortality[dwPos].dbDeadWolvesNatural += 1.0;
				} else if( pAliveDead[dwPos].cReason[d] == 'u' && bImputeUnknowns ) {
					if( dwPos / N_DAYS_IN_YEAR + 2004 <= 2011 ) {
						pMortality[dwPos].dbDeadWolvesHuman += IMPUTE_BEFORE_HUNT_HUMAN;
						pMortality[dwPos].dbDeadWolvesNatural += IMPUTE_BEFORE_HUNT_NATURAL;
					} else {
						pMortality[dwPos].dbDeadWolvesHuman += IMPUTE_AFTER_HUNT_HUMAN;
						pMortality[dwPos].dbDeadWolvesNatural += IMPUTE_AFTER_HUNT_NATURAL;
					}
				}
			}

		}
	}

	//consolidate all collected counts per year
	DWORD dwYear;
	for( DWORD dwPos = 0; dwPos < N_MAX_ENTRIES; dwPos++ ) {
		dwYear = dwPos / N_DAYS_IN_YEAR;
		if( pMortality[dwPos].dbDeadWolvesHuman > 0.0000001 ) {
			pMortalityOccured[dwYear].dbDeadWolvesHuman += pMortality[dwPos].dbDeadWolvesHuman;
		}
		if( pMortality[dwPos].dbDeadWolvesNatural > 0.0000001 ) {
			pMortalityOccured[dwYear].dbDeadWolvesNatural += pMortality[dwPos].dbDeadWolvesNatural;
		}
		pMortalityOccured[dwYear].dYear = dwYear + 2004;
	}

	if( cOnlySUMMARY ) { // if only summary required - consolidate in before and after hunt. The year entries are only two: 2011 means 7 years 2005-2011;  2012 means 7 years 2012-2018
		DWORD dwBeforeAfterHuntCount = 0;
		for( DWORD dwPos = 0; dwPos < N_YEARS; dwPos++ ) {  // we are discarding first and last year
			if( dwPos != 0 ) { // skip the first entry, as we are accumulating in entry 0 and 1
				if( pMortalityOccured[dwPos].dYear >= 2012 ) {
					dwBeforeAfterHuntCount = 1;
				}
				pMortalityOccured[dwBeforeAfterHuntCount].dbDeadWolvesHuman += pMortalityOccured[dwPos].dbDeadWolvesHuman;
				pMortalityOccured[dwBeforeAfterHuntCount].dbDeadWolvesNatural += pMortalityOccured[dwPos].dbDeadWolvesNatural;
			}
			if( dwPos == 1 ) { // and clear entry 1 first time, as we are going to be accumulating in it the second half of years
				pMortalityOccured[dwPos].dbDeadWolvesHuman = 0.0;
				pMortalityOccured[dwPos].dbDeadWolvesNatural = 0.0;
			} else {
				pMortalityOccured[dwBeforeAfterHuntCount].dYear = (dwBeforeAfterHuntCount == 0 ? 2011 : 2012);
			}
		}
		pMortalityOccured[2].dYear = 2019;
		pMortalityOccured[2].dbDeadWolvesHuman = pMortalityOccured[0].dbDeadWolvesHuman + pMortalityOccured[1].dbDeadWolvesHuman;
		pMortalityOccured[2].dbDeadWolvesNatural = pMortalityOccured[0].dbDeadWolvesNatural + pMortalityOccured[1].dbDeadWolvesNatural;
	}

	// produce the prefix to the file month with appropriate selections
	if( cSelectBySEX ) {   // if selecting by sex, do it only for adults, thus the age flag in file name is redundant
		cSelectByAGE = 0;  // also, if it wasn't specified, we may have added 'a' for age for entry selection to work properly
	}
	char szSelection1[2] = { cSelectByAGE,  '\0' };
	char szSelection2[2] = { cSelectBySEX, '\0'};
	// make the Dst file name
	_snprintf( szDstFile, sizeof( szDstFile ), "%s\\%s%s%s%s.csv", argv[2], szUnnownsImputation[bImputeUnknowns], szYearStart[nYearStartMonth], szSelection1, szSelection2);
	// open the Dst file
	HANDLE	hDstFile = CreateFile( szDstFile, GENERIC_READ | GENERIC_WRITE, FILE_SHARE_READ | FILE_SHARE_WRITE, NULL, CREATE_ALWAYS, FILE_ATTRIBUTE_NORMAL | FILE_FLAG_RANDOM_ACCESS, NULL );
	if( hDstFile == INVALID_HANDLE_VALUE ) {
		cout << "Failed to create destination file: " << szDstFile << "\n";
		return 0;
	}
	// create and write the title line, add Mortality only for summaries, as it is dome more precisely for individual years after the regression
	_snprintf( szTempBuf, sizeof( szTempBuf ), "Year,SurvivalRaw,MortalityRaw,RadioDays,Dead,DeadHuman,DeadNatural,DailyHazardVariance,DailyHazard,DailyHazardHuman,DailyHazardNatural%s\r\n", (cOnlySUMMARY?",MortalityHumanSmr,MortalityNaturalSmr,ciLwrSmr,ciUprSmr":"") );
	//	_snprintf( szTempBuf, sizeof( szTempBuf ), "Year,Mortality,Survival,RadioDays,Died\r\n");  // initial entries
	WriteFile( hDstFile, szTempBuf, (DWORD)strlen( szTempBuf ), &dwBytesWritten, NULL );

	DWORD dwCumulativeDays1 = 0;

	for( DWORD dwPos = 0; dwPos < N_YEARS; dwPos++ ) {
		if( cOnlySUMMARY && dwPos == 3 ) {
			break;
		}
		DWORD dWolfDays = 0;
		if( cOnlySUMMARY ) {
			DWORD dMaxYears = 8;
			DWORD dPosOffset = dwPos * 8;
			if( dwPos == 2 ) { // accumulate all 14 years for the third entry
				dMaxYears = 16;
				dPosOffset = 0;
			}
			for( DWORD dYearPos = 0; dYearPos < dMaxYears; dYearPos++ ) {
				DWORD dYearCounter = (dYearPos + dPosOffset); 
				dWolfDays += CountAliveWolfDays( pAliveDead, dYearCounter * N_DAYS_IN_YEAR );
			}
			dwCumulativeDays1 += dWolfDays;
		} else {
			dWolfDays = CountAliveWolfDays( pAliveDead, dwPos * N_DAYS_IN_YEAR );
			dwCumulativeDays1 += dWolfDays;
		}
		double dbDeadWolvesAll = pMortalityOccured[dwPos].dbDeadWolvesHuman + pMortalityOccured[dwPos].dbDeadWolvesNatural;
		double dbWolfDays = (double)dWolfDays;
		double dbFiniteDailySurvival = CalcMaximumLikelihoodSurvivalRate( dbWolfDays, dbDeadWolvesAll );
		double dbFiniteDailyHzd = 1.0 - dbFiniteDailySurvival;
		double dbFiniteDailyHzdHuman = 1.0 - ((dbWolfDays - pMortalityOccured[dwPos].dbDeadWolvesHuman) / dbWolfDays);
		double dbFiniteDailyHzdNatural = 1.0 - ((dbWolfDays - pMortalityOccured[dwPos].dbDeadWolvesNatural) / dbWolfDays);
		double dbSurvival = pow( dbFiniteDailySurvival, 365.0 );
		double dbUpper = 0.0, dbLower=0.0;

		double dbVariance = CalcYearlySurvConfIntervals( dbFiniteDailySurvival, dbWolfDays, dbDeadWolvesAll, &dbLower, &dbUpper );

		char szAssignedMortality[128] = "";
		if( cOnlySUMMARY ) {
			double dbMortalityHuman = 0.0;
			double dbMortalityNatural = 0.0;
			if( dbFiniteDailyHzd > 0.0000001 ) {
				dbMortalityHuman = (1.0 - dbSurvival) * dbFiniteDailyHzdHuman / dbFiniteDailyHzd;
				dbMortalityNatural = (1.0 - dbSurvival) * dbFiniteDailyHzdNatural / dbFiniteDailyHzd;
			}
			_snprintf( szAssignedMortality, sizeof( szAssignedMortality ), ",%.8g,%.8g,%.8g,%.8g", dbMortalityHuman, dbMortalityNatural, dbLower, dbUpper );
		}
		
		_snprintf( szTempBuf, sizeof( szTempBuf ), "%u,%.8g,%.8g,%u,%.8g,%.8g,%.8g,%.8g,%.8g,%.8g,%.8g%s\r\n", pMortalityOccured[dwPos].dYear, dbSurvival,
			1.0 - dbSurvival, dWolfDays,dbDeadWolvesAll, 
			pMortalityOccured[dwPos].dbDeadWolvesHuman, pMortalityOccured[dwPos].dbDeadWolvesNatural,
			dbVariance, dbFiniteDailyHzd, dbFiniteDailyHzdHuman, dbFiniteDailyHzdNatural,
			szAssignedMortality);
		WriteFile( hDstFile, szTempBuf, (DWORD)strlen( szTempBuf ), &dwBytesWritten, NULL );
	}

	dwCumulativeDays1 = dwCumulativeDays1;

	CloseHandle( hDstFile );

    return 0;
}


BOOL ReadFileInMemBuffer( LPSTR szArgvFileName, BYTE **	ppbFileBfr, __int64 * piFileSize )
{
	char szSrcFile[MAX_PATH];
	
	_snprintf( szSrcFile, sizeof( szSrcFile ), "%s", szArgvFileName );
	HANDLE	hSrcFile = CreateFile( szSrcFile, GENERIC_READ | GENERIC_WRITE, FILE_SHARE_READ | FILE_SHARE_WRITE, NULL, OPEN_EXISTING, FILE_ATTRIBUTE_NORMAL | FILE_FLAG_RANDOM_ACCESS, NULL );
	if( hSrcFile == INVALID_HANDLE_VALUE ) {
		cout << "Failed to open file: " << szSrcFile << "\n";
		return FALSE;
	}


	LARGE_INTEGER liFileSize;
	GetFileSizeEx( hSrcFile, &liFileSize );

	BYTE *	pbFileBfr = (BYTE *)GlobalAllocPtr( GPTR, liFileSize.QuadPart );
	if( pbFileBfr == NULL ) {
		cout << "Failed to allocate memory for file: " << szSrcFile << "\n";
		CloseHandle( hSrcFile );
		return FALSE;
	}


	// read the whole file in the buffer, because the DWORD limit, reading it in quarter DWORDS - 1GB chunks, so we don't over-abuse the system buffering
	__int64 iBytesPos = 0;
	DWORD dReadBytesChunk = 0x400000;
	do {
		if( (__int64)dReadBytesChunk > (liFileSize.QuadPart - iBytesPos) ) {
			dReadBytesChunk = (DWORD)(liFileSize.QuadPart - iBytesPos);
		}

		DWORD dwBytesRead;
		if( !ReadFile( hSrcFile, &pbFileBfr[iBytesPos], dReadBytesChunk, &dwBytesRead, NULL ) || dReadBytesChunk != dwBytesRead ) {
			cout << "Failed to read file: " << szSrcFile << "\n";
			GlobalFreePtr( pbFileBfr );
			CloseHandle( hSrcFile );
			return FALSE;
		}
		iBytesPos += (__int64)dReadBytesChunk;
	} while( iBytesPos < liFileSize.QuadPart );

	*piFileSize = iBytesPos;
	*ppbFileBfr = pbFileBfr;

	CloseHandle( hSrcFile );

	return TRUE;
}


__inline BOOL GetTextLine( BYTE * pbFileBfr, __int64 * piBufPos, char * szTempBuf, int nTempBufSize, __int64 iFileSize )
{
	__int64 iBufPos = *piBufPos;
	__int64 iStartPos;

	// discard new lines
	while( (pbFileBfr[iBufPos] == '\r' || pbFileBfr[iBufPos] == '\n') && iBufPos < iFileSize ) {
		iBufPos++;
	}
	iStartPos = iBufPos;
	if( iBufPos >= iFileSize ) {
		return FALSE;
	}

	while( (pbFileBfr[iBufPos] != '\r' && pbFileBfr[iBufPos] != '\n') && iBufPos < iFileSize) {
		iBufPos++;
	}

	if( iBufPos >= iFileSize && iBufPos - iStartPos < 1) {  // finished reading file; if no data to load (at least 4 bytes) - done
		return FALSE; // can do no more
	}

	if( szTempBuf ) {
		int nLen = (int)min( nTempBufSize - 1, iBufPos - iStartPos + 1 );
		szTempBuf[nLen] = 0;

		lstrcpynA( szTempBuf, (char *)&pbFileBfr[iStartPos], nLen );
	}
	*piBufPos = iBufPos;

	return TRUE;
}

DWORD CountTextLines( BYTE * pbFileBfr, __int64 iFileSize )
{
	__int64 iBufPos = 0;
	DWORD dwLines = 0;

	// discard new lines
	while( GetTextLine( pbFileBfr, &iBufPos, NULL, 0, iFileSize ) ) {
		dwLines++;
	}

	return dwLines;
}


void CreateDir(char * szPath)
{
	int i= 0,j,k= 0;
	char szTempBuff[MAX_PATH];

	j= lstrlen((LPSTR)szPath);
	strncpy((LPSTR)szTempBuff, (LPSTR)szPath, sizeof(szTempBuff));
	for( i= i; i<=j; i++ ) {
		if( szTempBuff[i]=='\\' || szTempBuff[i]=='\0' ) {
			if( k==0 ) {
				k++;
			} else {
				szTempBuff[i]= '\0';
				_mkdir(szTempBuff);
				lstrcpy((LPSTR)szTempBuff, (LPSTR)szPath);
			}
		}
	}
}

double TrentRongstad_DailySurvivalEst( double dbWolfDays, double dbDeadCount ) // Trent and Rongstad (1974)
{
	return (dbWolfDays - dbDeadCount) / dbWolfDays;
}

double CalcMaximumLikelihoodSurvivalRate( double dbWolfDays, double dbDeadCount )
{
	if( dbDeadCount < 0.0000000001 ) {
		return 1.0;
	}
	double dbInitialSurv = TrentRongstad_DailySurvivalEst( dbWolfDays, dbDeadCount );
	double dbFinalSurv = 0.0, dbDiff=0.0;
	int nIterations = 0;
	do {
		double dbA = ((double)1.0 / dbInitialSurv) * (dbWolfDays - dbDeadCount - (dbDeadCount * dbInitialSurv) / (1.0 - dbInitialSurv));
		double dbB = ((double)1.0 / pow( dbInitialSurv, 2.0 )) * (dbWolfDays - dbDeadCount + (dbDeadCount * pow( dbInitialSurv, 2.0 )) / pow( (1.0 - dbInitialSurv), 2.0 ));
		dbFinalSurv = dbInitialSurv + dbA / dbB;
		dbDiff = dbFinalSurv - dbInitialSurv;
		nIterations++;
		dbInitialSurv = dbFinalSurv;
	} while( abs( dbDiff ) > 0.0000000001 );
	return dbFinalSurv;
}

double CalcStdErr( double dbWolfDays, double dbDeadCount )
{
	double dbStdError = pow( ((dbWolfDays - dbDeadCount) * dbDeadCount) / pow( dbWolfDays, 3.0 ), 0.5 );
	return dbStdError;
}

double CalcYearlySurvConfIntervals( double dbDailySurv,  double dbWolfDays, double dbDeadCount, double * pdbLower, double * pdbUpper )
{
	double dbStdError = CalcStdErr( dbWolfDays, dbDeadCount );
	//if( dbDeadCount == 0.0 ) { 
	//	dbStdError = CalcStdErr( dbWolfDays, 1 );
	//}
	double db95pc = 1.96 * dbStdError;
	if( dbDailySurv - db95pc < 0.0 ) {
		*pdbLower = 0.0;
	} else {
		*pdbLower = pow( dbDailySurv - db95pc, 365 );
	}
	*pdbUpper = pow( dbDailySurv + db95pc, 365 );
	double dbYearlySurvival = pow( dbDailySurv, 365 ); // already have it, just for debugging purposes here

	return pow(dbStdError, 2.0); // return variance
}
```
